# Parasitism by, species composition, morphometry, and parasitoidism of *Protocalliphora* bird blowflies (Diptera: Calliphoridae) in Quebec’s farmlands: a 16-year dataset

**DOI:** 10.1101/2025.07.23.666426

**Authors:** Simon Coroller-Chouraki, Jade Savage, Fanie Pelletier, Dany Garant, Marc Bélisle

## Abstract

Bird blowflies, *Protocalliphora* spp. (Diptera: Calliphoridae), are prevalent ectoparasites of altricial bird nestlings across the Holarctic region. Yet, their spatial and temporal dynamics of infestations, species composition, and interactions with parasitoids remain poorly understood. We present a 16-year (2004–2019) multisite study of bird blowfly infestations based on 2673 tree swallow, *Tachycineta bicolor* (Vieillot) (Passeriformes: Hirundinidae), nests collected across a 10 200-km² gradient of agricultural intensity in Quebec, Canada. Nest infestation prevalence and parasitic load varied markedly across space and time but showed synchronous recurrence at approximately 75% of sites, suggesting the influence of regional and local processes. Yearly rates of parasitoidism of bird blowfly puparia by *Nasonia* spp. wasps (Hymenoptera: Pteromalidae) were high but variable (48–90%), likely contributing to the temporal fluctuations in bird blowfly prevalence and load. Substantial interannual shifts in the relative abundance of *Protocalliphora* species (*P. bennetti*, *P. metallica*, and *P. sialia*) emphasised the importance of species-level identification in bird blowfly ecological studies. Large overlap in puparia size among species challenged the utility of traditional diagnostic traits for species identification. Finally, dormancy or mortality of *Nasonia* spp. occurred in 3–16% of *Protocalliphora* spp. puparia, depending on year. These findings highlight the importance of long-term, multitrophic, and spatially explicit monitoring to unravel the drivers of host–parasite–parasitoid dynamics.

## Introduction

Avian ectoparasites encompass an exceptionally diverse array of invertebrate taxa, ranging from leeches (Hirudinea) and ticks (Ixodida) to insects from various orders, including lice (Phthiraptera), true bugs (Hemiptera), fleas (Siphonaptera), and flies (Diptera) (Rothschild and Clay 1957; Clayton *et al*. 2010). Many of these ectoparasites are hematophagous, a feeding mode that has evolved independently multiple times within insects, including in Diptera, (Deepak and Shruti 2024; Cardoso *et al*. 2025), and which occurs in many families of both the Nematocera and Brachycera suborders (*e.g*., Culicidae, Simuliidae, Ceratopogonidae, Hippoboscidae, Nycteribiidae). However, significant differences exist among dipteran ectoparasites in terms of host selection and specificity, whether they are obligatory blood feeders or not, and regarding diet differentiation across life stages. For instance, the larvae of *Protocalliphora* spp. (Diptera: Calliphoridae), also known as bird blowflies, are obligatory blood-feeding parasites of nestlings belonging to a wide range of altricial bird species. Little is known of the flies’ adult diet, but they have been known to visit the flowers of herbaceous plants (Sabrosky *et al*. 1989; Bennett and Whitworth 1991). Although the biology and ecology of hematophagous flies parasitising humans and domestic animals are well known (*e.g*., Marquardt *et al*. 2004), flies parasitising wild animals have received less attention or have mainly been studied according to the costs they can inflict on their hosts. Bird blowflies exemplify this case, with most previous studies focusing on measuring the impacts their larvae may have on the nestlings and reproductive success of altricial birds (Simon *et al*. 2004; Maziarz *et al*. 2022). In recent years, however, several studies have addressed the environmental factors that drive the prevalence and intensity of bird nest infestations by *Protocalliphora* spp. (Mennerat *et al*. 2021; Merino *et al*. 2025; González-Bernardo *et al*. 2025). However, only a few studies to date have formally investigated the determinants of bird nest infestations by these parasites across multiple years and sites (Mennerat *et al*. 2021; Maziarz *et al*. 2022; Merino *et al*. 2025; González-Bernardo *et al*. 2025). Moreover, these studies, which mainly focused on the effects of weather or climatic factors, were conducted in a small number of forested sites and, therefore, on a limited range of habitat heterogeneity. Because of this, we have limited information on the spatial and temporal variability of infestations and therefore on the influence of abiotic and biotic factors on the host–parasite dynamics involving *Protocalliphora* species. The lack of long-term monitoring data on bird nest infestations by these parasites also concerns the species of blowflies involved in these interactions. At least 40 species of *Protocalliphora* have been described in the Holarctic region (Sabrosky *et al*. 1989), many of which have overlapping distribution ranges, resulting in occasional mixed infestations within a bird nest (Daoust *et al*. 2012a). Moreover, the morphological identification of species is challenging in this group, at least for nonspecialists, and has been considered to be reliable only in adults (Heeb *et al*. 2000). Furthermore, locating and sampling adult bird blowflies in the field is not easy (Stiner 1969; Bennett and Whitworth 1991), and larvae and puparia often remain the primary evidence of bird blowfly parasitism and parasitic load (Jánošková *et al*. 2010). Despite significant progress regarding the taxonomy, phylogeny (Kutty *et al*. 2019; Yan *et al*. 2021), and identification of *Protocalliphora* species, notably through the work of Sabrosky *et al*. (1989) and Whitworth (2002, 2003, 2019), identification based on puparia morphology is complicated by the fact that key criteria, such as distance between specific spines, prothoracic fringe diameter, and distance between posterior spiracles, can vary widely within and overlap between species (Jánošková *et al*. 2010; Jamriška and Modrý 2013). Furthermore, identification based on DNA barcoding using the Folmer region of the common *CO*1 gene fails to discriminate between most described *Protocalliphora* species (Whitworth *et al*. 2007). These identification challenges, especially when considering the high parasite load found in some nests, may have led some ecologists to neglect bird blowfly species composition and typically lump all individuals under “*Protocalliphora* spp.” (*e.g.*, Heeb *et al*. 2000; Bańbura *et al*. 2004; Hannam 2006; Musgrave *et al*. 2019). Lumping all species together can, however, bias species abundance estimates and hinder the study of host–parasite dynamics by neglecting the potential competition that may occur among *Protocalliphora* species, as well as their interactions with parasitoids within host nests.

Other overlooked factors that may influence parasitoid–parasite interactions include inter-and intraspecific variation in body size. Such variation may reflect differences in larval growth conditions, which in turn can affect metamorphosis, emergence success, and susceptibility to host-selection by gregarious jewel wasps, *Nasonia* spp. (Hymenoptera: Pteromalidae) (Werren and Loehlin 2009). These generalist wasps, which primarily parasitise the pupal stage of *Protocalliphora* spp., are also reported to parasitise several other calyptrate Diptera pupae of agricultural relevance (Rueda and Axtell 1985), thereby potentially contributing to the top–down regulation of these flies. Most research on *Nasonia* spp., including the well-known *N. vitripennis* (Walker) and *N. giraulti* Darling, which both attack *Protocalliphora* spp. (Daoust *et al*. 2012a; Garrido-Bautista *et al*. 2020, 2024), have largely focused on their genetics and behavioural ecology under laboratory conditions (*e.g*., Werren and Loehlin 2009; Deveux *et al*. 2023). Past studies therefore provide limited insights into the ecological interactions of this parasitoid–parasite relationship in natural habitats and into how the parasitoid’s top–down regulation potential against bird blowflies fluctuates under different conditions in either space or time (Garrido-Bautista *et al*. 2024). These gaps in our understanding of *Protocalliphora* spp. extend beyond taxonomy, morphometry, and distribution. Long-term studies focusing on the prevalence and abundance of bird blowfly larvae within nests have noted important annual fluctuations after accounting for some environmental conditions (Wesołowski 2001; Musgrave *et al*. 2019; Merino *et al*. 2025); however, only Merino *et al*. (2025) reported patterns suggesting cyclical abundance, although they did not formally analyse cycles. To our knowledge, no details on temporal variation in species composition and parasitoidic pressures have been documented. In the present study, we quantify the spatial and temporal variability in the prevalence, intensity, and species composition of tree swallow, *Tachycineta bicolor* (Vieillot) (Passeriformes: Hirundinidae), nest infestations by bird blowflies over a 16-year period within a 10 200-km² gradient of agricultural intensity in southern Quebec, Canada. We also report species-specific size distributions for the puparia of three species of *Protocalliphora* and temporal variation in parasitoidism rates by *Nasonia* spp., as well as the rate of parasitoid dormancy (*i.e.,* winter diapause) within the host’s puparia.

## Methods

### Study system and field sampling

We collected *Protocalliphora* spp. puparia between 2004 and 2019 within a network of 400 nest boxes distributed equally among 40 farms located along a 10 200-km² gradient of agricultural intensity in southern Quebec, Canada (Ghilain and Bélisle 2008; Supplementary material, File S1). Nest boxes were placed about 1.5 m above ground and spaced by about 50 m apart along linear transects that followed field edges. Although the primary objective of the above network initiated in 2004 was to quantify the impact of land use on tree swallow prey availability (Powell *et al*. 2024) and breeding ecology (Garrett *et al*. 2022a), it evolved into a comprehensive platform for diverse investigations, including the effects of bird blowfly parasitic load on the swallows’ breeding success (Daoust *et al*. 2012b; Pigeon *et al*. 2013; Sigouin *et al*. 2021). At the end of each breeding season, we collected and bagged the entire content of nest boxes exclusively occupied by tree swallows, a single-brooded species. Collected nests were stored at 4 °C prior to processing, which included 2 days at –80 °C to kill any remaining living organisms, before sorting under a ventilated hood. *Protocalliphora* spp. puparia found within nest material were counted and stored in vials filled with 75% ethanol and referenced according to farm and nest box IDs and year of sampling. Note that puparia from 2008 and 2009 were collected under two protocols by Daoust *et al*. (2012a) to rear some material in the lab and to collect emerging flies or *Nasonia* spp. wasps. As a result, some specimens for these two years were no longer available for further examination in the present study (*i.e.*, for parasitoidism detection or for bird blowfly species composition analyses and morphometry). Nonetheless, our dataset allowed us to investigate nest infestation and parasitic load using nests from all nest boxes occupied by tree swallows with a single clutch event that resulted in at least one nestling on a given year between 2004 and 2019 (including 2008 and 2009), for a total of 2673 nests (Table 1).

**Table 1.** Number of tree swallow nests and *Protocalliphora* spp. puparia collected and analyzed between 2004 and 2019 in southern Quebec, Canada. No puparia from 2008 to 2009 could be included in our parasitoidism, species composition, and morphometry estimates because they were used in a previous study by Daoust *et al*. (2012a) and were no longer suitable for further processing. Instead, proportions for 2008 and 2009 were directly derived from Daoust *et al*. (2012a).

| Samples (#) | Datasets |  |  |  |
| --- | --- | --- | --- | --- |
|  | All data | Parasitoidism | Species composition | Morphometry |
| Nests examined | 2673 | 263 | 263 | 263 |
| Nests infested by *Protocalliphora* spp. with > 1 puparium suitable for processing | 1227 (45.90%) | 259 | 216 | 174 |
| Nests infested by *Nasonia* spp. | — | 218 | — | — |
| Puparia examined | 22 630 | 3994 | 3994 | 3994 |
| Puparia excluded (damaged) | — | 386 | 985 | 2012 |
| Puparia retained | — | 3608 | 3009 | 1982 |
| Puparia without *Nasonia* spp. | — | 1125 (31.18%) | — | — |
| Puparia with *Nasonia* spp. | — | 2483 (68.82%) | — | — |
| Puparia containing nonemerged *Nasonia* spp. | — | 318 | — | — |
| Puparia of *P. sialia* | — | — | 2426 | 1569 |
| Puparia of *P. bennetti* | — | — | 474 | 347 |
| Puparia of *P. metallica* | — | — | 109 | 66 |

### Specimen examination

Due to the considerable number of *Protocalliphora* spp. puparia and the significant time required for individual specimen examination, a subsample of 263 nests infested by *Protocalliphora* spp. was selected for parasitoidism assessment, species identification, and morphometric measurements of puparia found therein (Table 1). Infested nests were subsampled as follows: 10 farms were selected by stratified random sampling to represent the gradient of agricultural intensity of the study area across years. (For details, see Supplementary material, File S2.) For each year (excluding 2008 and 2009), two infested nests (when available) were randomly sampled on each of the above 10 farms. All puparia found within these nests were retained. If fewer than 100 puparia could be obtained for a given year, we randomly selected an additional farm from which we randomly selected up to two nests to extract all puparia therein. This process was then repeated until at least 100 puparia could be processed for a given year. The number of additional farms thereby sampled averaged 1.2 per year, a number unlikely to alter the general patterns provided by the core group of 10 farms.

All *Protocalliphora* spp. specimens from the resulting subsample of 263 nests (n = 3994 puparia) were examined, and only those with adequate structural integrity for the determination or measurement of a given attribute were considered in data analyses regarding parasitoidism, species composition, and morphometry (Table 1).

### Parasitoidism of *Protocalliphora* spp. by *Nasonia* spp

Species-level identification of *Nasonia* spp. wasps was not conducted in the present work due to fall sample collections, when most parasitoids had already emerged or when those that had not were killed before completing their development. However, early subsampling and rearing of *Protocalliphora* spp. pupae by Daoust *et al*. (2012a) within the same study system allowed the post-emergence identification of two *Nasonia* species — *N. vitripennis* and *N. giraulti*. *Nasonia vitripennis* accounted for more than 90% of parasitoidal events in *Protocalliphora* spp.

The occurrence of *Nasonia* spp. parasitoidism on a host puparium is typically diagnosed by the presence of one or more exit holes of 0.5–1 mm in diameter. Although other parasitoids can create similar emergence holes, *N. vitripennis* and *N. giraulti* were the only two parasitoid species to emerge from 956 pupae sampled by Daoust *et al*. (2012a) across the same study system in 2008–2009. We therefore presume that most, if not all, emergence holes noted in sampled *Protocalliphora* spp. resulted from individuals of this genus. Because *Nasonia* spp. can die or overwinter as pupae inside their host instead of emerging at the end of the active season (Floate and Skovgård 2004; Garrido-Bautista *et al*. 2020), puparia containing a dead *Protocalliphora* spp. pupa and those containing unemerged *Nasonia* spp. look identical. For this reason, we systematically dissected *Protocalliphora* spp. puparia showing no sign of either fly or wasp emergence to verify the occurrence of *Nasonia* spp. parasitoids to discriminate pupation failures linked to parasitoidism from those resulting from other causes. All 3,608 *Protocalliphora* spp. puparia assessed for *Nasonia* spp. infestation were therefore classified into one of the following categories: no evidence of *Nasonia* spp. parasitoidism; presence of *Nasonia* spp. emergence holes; and presence of unemerged *Nasonia* spp. pupae without exit holes.

### *Protocalliphora* species composition

We identified 3009 well-preserved *Protocalliphora* spp. puparia based on morphology, following Sabrosky *et al*. (1989) and Whitworth (2002, 2003, 2019), with additional feedback from T.L. Whitworth (personal communication). Consistently with Daoust *et al*. (2012a), who conducted research within the same study system, only three species were identified: *Protocalliphora sialia* Shannon and Dobroscky, *Protocalliphora bennetti* (Whitworth), and *Protocalliphora metallica* (Townsend). Ten puparia (five with and five without *Nasonia* exit holes) from each of the three *Protocalliphora* species were kept as voucher specimens and deposited in the Bishop’s University Insect Collection (BUIC; Sherbrooke, Quebec, Canada), along with the remains of five dissected specimens containing unmerged parasitoid wasps. All material is filed in the voucher collection under “Coroller-Chouraki *Protocalliphora* spp. 2025”.

### *Protocalliphora* spp. morphometry

To assess inter- and intraspecific variation in the maximum length and width of *Protocalliphora* spp. puparia (in millimetres), each puparium was photographed before species identification using a Leica M165FC stereomicroscope equipped with a 0.63× planApo objective, a Leica MC170HD colour camera, and Leica LAS× image-capture software, version 3.0.3. The microscope being already factory-encoded, all acquired images were automatically calibrated in terms of pixel size (µm/pixel). All image-based measurements were subsequently performed using Fiji image analysis software (Schindelin *et al*. 2012). Image analysis was automated using a homemade Fiji script developed using the built-in Fiji/ImageJ API. For each image, a threshold-based image segmentation was performed to isolate individual puparia, and a *Fit Ellipse* function was applied to each object. This allowed us to model any photographed puparium as an ellipsoid-shaped object and to calculate its theoretical maximum length and width. This approach minimised measurement biases that might result from puparia from which an adult fly had emerged otherwise appearing shorter due to the missing emergence cap.

### Data analysis

Interannual variability in tree swallow nest infestation by *Protocalliphora* spp. between 2004 and 2019 was evaluated by calculating, for each sampling year, the proportion of infested nests, and the average number of *Protocalliphora* spp. puparia per infested nest (± standard deviation). We then assessed how the annual mean nest infestation rate across farms varied in time between 2004 and 2019 and to what level each farm followed that global trend, if any, using a generalised linear additive mixed model fitted with the *mgcv* package, version 1.9-1 (Wood 2023) in R, version 4.3.1 (R Core Team 2023). More specifically, we modelled the proportion of infested nests over time based on two components: (1) a global smooth term for *year* to capture the overall temporal trend, and (2) farm-specific smooth terms to account for temporal patterns unique to each farm. The farm-specific (local) trends could differ in shape and complexity from the global one (see Pedersen *et al*. 2019, Model GI). The generalised linear additive mixed model was fitted using a binomial distribution on farm–year counts of infested nests (with the farm–year nest counts as weight argument) and logit-link function. We performed the same exploratory analyses for the average number of *Protocalliphora* spp. puparia per infested nest with a generalised linear additive mixed model that used a tweedie distribution and a log-link function due to substantial overdispersion with respect to a (truncated) Poisson distribution. Temporal autocorrelation in the residuals was estimated to be weak (*r_k_* < 0.15) at all *k* (≤ 10) time lags using the *acf* function in R (Simpson 2018), suggesting that the fitted generalised linear additive mixed models adequately captured the temporal structure within the response variables. Detailed codes, model diagnostics, and results for these analyses are provided in Supplementary material, File S3.

Interannual variability in parasitoid–parasite dynamics was evaluated by contrasting the yearly proportion of noninfested to infested puparia and the proportion of infested puparia from which wasps had emerged against those still containing *Nasonia* spp. pupae or adults. Interannual variability in *Protocalliphora* species composition was assessed by calculating the yearly relative proportion of each species for all specimens that could be unambiguously identified. Parasitoidism rates and species composition for 2008–2009 were obtained from Daoust *et al*. (2012a).

Intra- and interspecies variability in the maximum length and width of puparia of *P. sialia*, *P. bennetti*, and *P. metallica* were assessed first by using density plots and boxplots based on the raw data. We then assessed interspecific differences in mean puparium maximum length and width using generalised linear mixed models fitted with the *glmmTMB* R package (Brooks *et al*. 2023). In this analysis, *P. bennetti*, the intermediate-size species (Sabrosky *et al*. 1989; Whitworth 2002), was designated as the reference category. *Farm ID* and *year* were included as random effects to account for environmental variation and to control for potential confounding effects. *Nest box ID* was included as a random effect, but this led to convergence issues due to the low number of repeated measures within each nest: for this reason, this variable was omitted from the analysis. The proportion of variance explained by *Farm ID* and *year* random effects were estimated following Nakagawa and Schielzeth (2013). Species-specific maximum length and width being approximately normally distributed, generalised linear mixed models were fitted using a Gaussian distribution and identity link.

## Results

### Spatiotemporal variation in *Protocalliphora* spp. prevalence and load

Nest infestation by bird blowflies occurred widely throughout the study area. All 40 monitored farms harboured at least one infested tree swallow nest at some point between 2004 and 2019 and between 67.65% and 92.31% of farms had at least one infested nest, depending on the year. Overall, 45.90% (1227) of the 2673 tree swallow nests with a single clutch event leading to at least one nestling were infested by at least one *Protocalliphora* spp. larva, based on counts of puparia. The proportion of infested nests varied substantially both among years (range: 28.76% in 2012 to 69.33% in 2008; Fig. 1A) and among farms (range: 13.37–79.49%). Although most farms showed yearly fluctuations in nest infestation rate that followed the global, nonlinear trend suggestive of a recurrent pattern with an approximately 3-year period at the scale of the study area, about 30% of farms showed mild to strong departures from that global trend (n = 587 farm–years; GAMM *R*^2^= 48.8%; Supplementary material, File S3).

**Figure 1.**
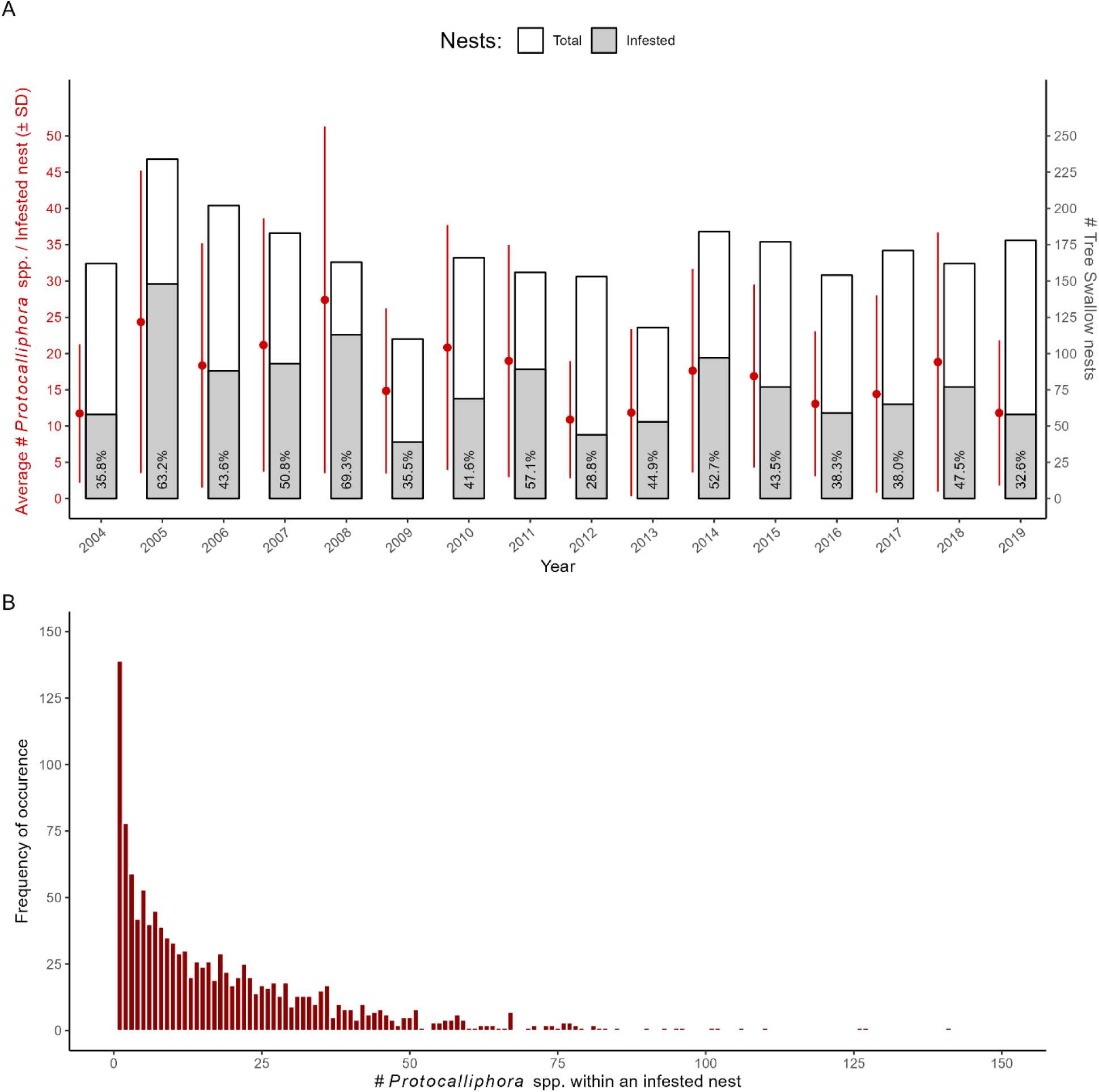
**A**, Patterns of tree swallow nest infestation by *Protocalliphora* spp. observed between 2004 and 2019 in southern Quebec, Canada. Left y-axis and corresponding point ranges (red dots ± error bars) show the yearly average (± standard deviation) number of *Protocalliphora* spp. puparia per infested nest. Right y-axis and corresponding stacked bar graph show the yearly number of infested nests (grey) over the yearly total number of examined nests (*i.e.,* nests that harboured a single clutch that experienced at least one hatching event; white). **B**, Histogram showing the frequency distribution of the number of *Protocalliphora* spp. puparia found within infested nests (*n* = 1227).

A total of 22 630 puparia were counted among infested nests, resulting in an overall estimated average load (± standard deviation) of 18.44 ± 19.16 *Protocalliphora* spp. per infested nest. The average number of *Protocalliphora* spp. per infested nest exhibited variability both within and across years, fluctuating between 10.89 ± 8.08 (2012) and 27.40 ± 23.90 (2008) with a yearly coefficient of variation ranging between 74.21 and 97.11% (Fig. 1A). The overall frequency distribution of the number of *Protocalliphora* spp. per infested nest was, however, highly skewed to the right with a median of 12 puparia (range: 1–141 puparia; Fig. 1B). Interestingly, most farms showed yearly fluctuations in average *Protocalliphora* spp. loads per infested nest that followed a global, nonlinear trend, which peaks and troughs coincided with those observed for the proportion of nests infested by *Protocalliphora* spp. (n = 1227 infested nests; GAMM *R*^2^ = 12.2%; Supplementary Material 3). However, the farms that deviated from the global load trend (*i.e.*, 22.5% of farms) were not the same as those for which the proportion of infested nests and the generalised linear additive mixed model residuals showed no temporal autocorrelation pattern.

### Parasitoidism of *Protocalliphora* spp. by *Nasonia* spp

*Nasonia* spp. likely occurred throughout the study area because they were detected at least once in 30 of the 31 farms from which *Protocalliphora* spp. puparia were subsampled to estimate parasitoidism rate. Overall, *Nasonia* spp. parasitised at least one puparium in 85.08% of nests in which *Protocalliphora* spp. were found and 68.82% of the 3608 subsampled puparia that could be examined (Table 1). Importantly, 12.81% of parasitised *Protocalliphora* spp. puparia contained nonemerged *Nasonia* spp. individuals. These nonemerged individuals occurred nearly exclusively at the pupal stage, suggesting that they either had failed to produce adults or had entered diapause for the coming winter. Parasitoidism by *Nasonia* spp. remained high across years, ranging from 48.25% (2006) to 90.48% (2012) of *Protocalliphora* spp. puparia (Fig. 2). Note that Daoust *et al*. (2012a) estimated parasitoidism rates in 2008 and 2009 based on a mix of reared *Protocalliphora* spp. pupae and on the presence of exit holes made by *Nasonia* spp. to exit puparia. They reported among the lowest parasitoidism rates of the study period (*i.e*., 50.35% and 39.29%, respectively). The proportions of infested *Protocalliphora* spp. puparia from which *Nasonia* spp. did not emerge before winter varied substantially, ranging from 3.07% in 2006 to 16.36% in 2019 (Fig. 2).

**Figure 2.**
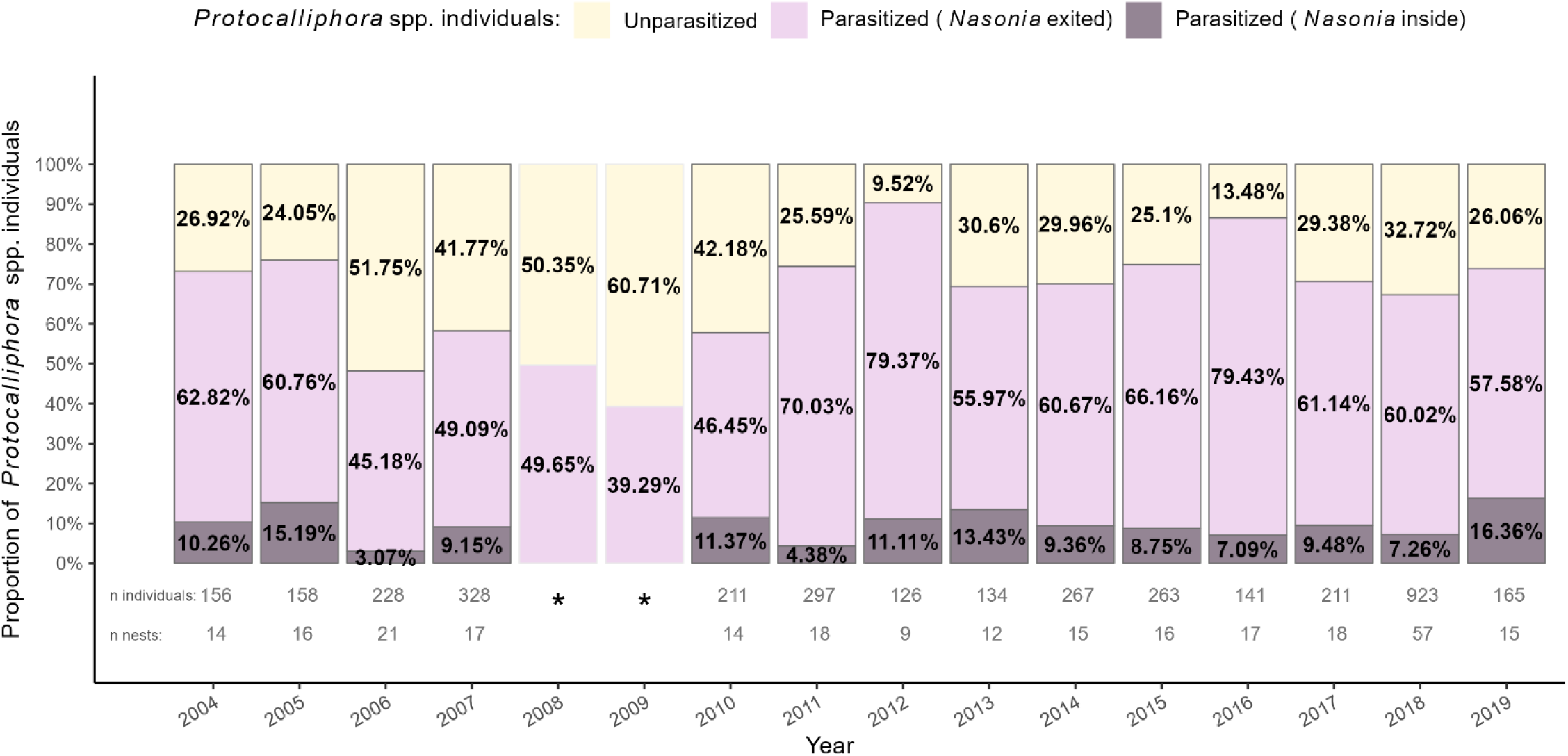
Stacked bar graph representing the interannual rates of *Protocalliphora* spp. puparia parasitoidism by *Nasonia* spp. wthin tree swallow nests collected in southern Quebec, Canada, between 2004 and 2019. *Protocalliphora* spp. puparia were classified as unparasitised when they showed no signs of parasitoidism (yellow) and as parasitised if *Nasonia* spp. exit holes were observed on the puparia (pink) or if *Nasonia* spp. first instars were found when *Nasonia* spp. exited *Protocalliphora* spp. (purple). *Proportions for 2008 and 2009 were calculated from statistics reported by Daoust *et al*. (2012a, tables 3 and 4). Parasitoidism was determined by Daoust *et al*. (2012a) using a mix of reared *Protocalliphora* spp. pupae and based on the presence of holes made by *Nasonia* spp. to exit puparia.

### *Protocalliphora* species composition

Of the 3009 *Protocalliphora* spp. puparia suitable for species identification, 82.62% belonged to *P. sialia*, 15.75% belonged to *P. bennetti*, and 3.62% belonged to *P. metallica* (Table 1). *Protocalliphora sialia* was consistently the most abundant species in each year, ranging from 61.72% in 2011 to 98.41% the following year (Fig. 3). In contrast, *P. bennetti* exhibited more pronounced fluctuations in relative abundance, ranging from quasi-absence (0.79%) in 2012 to 32.01% in 2011. *Protocalliphora metallica*, the least abundant species in our study system, also showed important variation in its relative abundance: it was either completely absent from our subsamples in 2005 and 2016 or as or more abundant than *P. bennetti* was in 2004, 2012, and 2017, but it never exceeded 9.27% (2017) of *Protocalliphora* puparia in any given year (Fig. 3). Data extrapolated from Daoust *et al*. (2012a) for 2008 and 2009 also showed a strong dominance of *P. sialia*, with proportions of 96.59% and 90.74%, respectively. Overall, all three species were found over most of the study area (Supplementary material, File S4).

**Figure 3.**
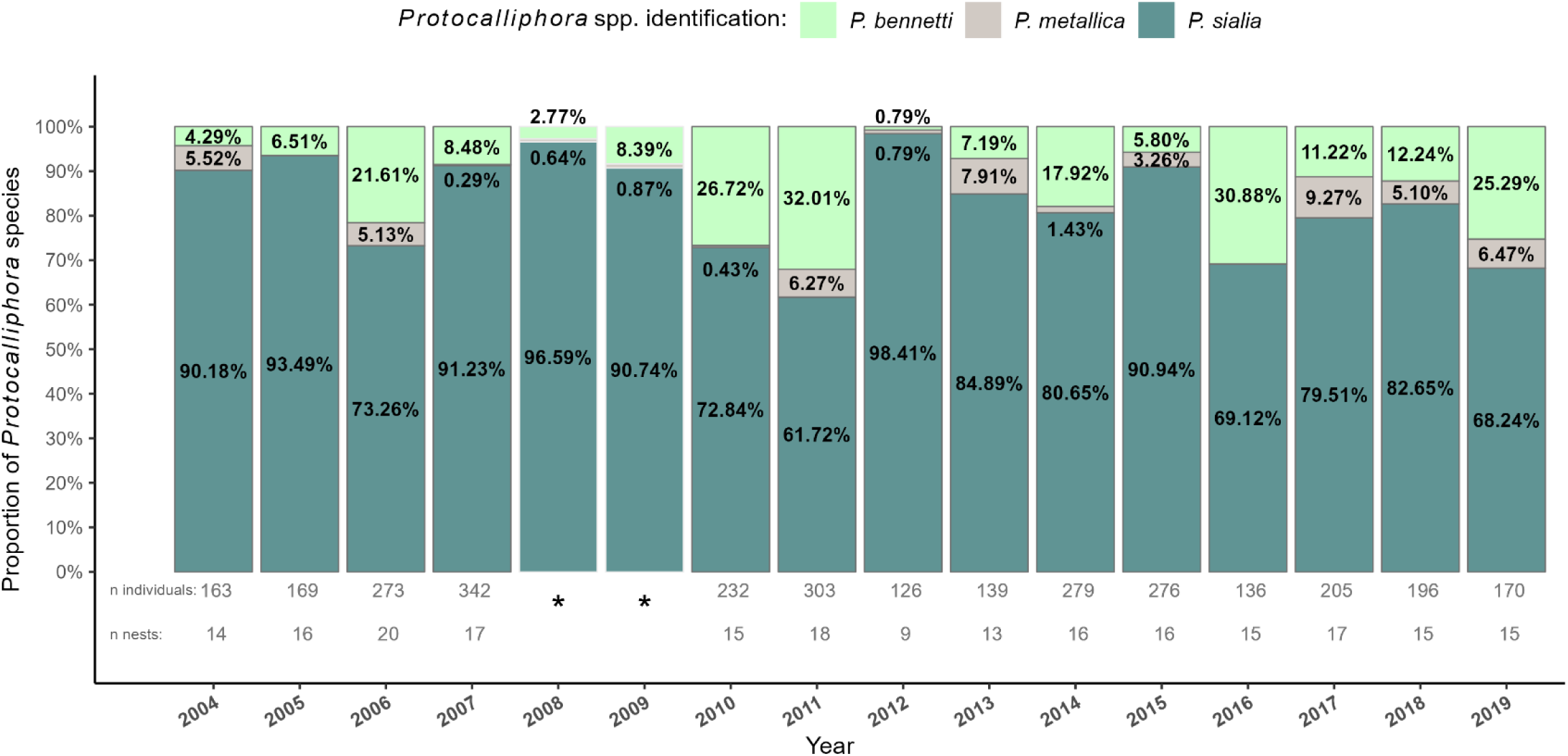
Stacked bar graph representing the interannual *Protocalliphora* species composition within tree swallow nests collectd in southern Quebec, Canada, between 2004 and 2019. Three species were identified (*P. bennetti*, light green; *P. metallica*, grey; *P. sialia*, dark green). *Proportions for 2008 and 2009 were derived from Daoust *et al*. (2012a; table 1).

### *Protocalliphora* spp. morphometry

The maximum puparium lengths and widths of the three *Protocalliphora* species were obtained from a subsample of 1982 individuals (Table 1). The distribution of maximum lengths and widths of the three species was unimodal and relatively symmetrical, although slightly skewed to the left in *P. metallica* (Fig. 4). Despite substantial intraspecific variation leading to distributional overlap, the generalised linear mixed models showed the species differed in their average morphometry. *Protocalliphora bennetti* exhibited a mean puparium length (± standard deviation) of 8.18 ± 0.67 mm (range: 5.24–9.93 mm), whereas *P. metallica* showed a significantly shorter mean length (7.34 ± 0.71 mm; range: 5.82–9.29 mm; *P* < 0.001) and *P. sialia* showed a longer one (8.74 ± 0.58 mm; range: 5.70–10.27 mm; *P* < 0.001). Regarding width, *P. metallica* had a narrower puparium (3.36 ± 0.41 mm, range: 2.25–4.16 mm; *P* < 0.001) and *P. sialia* a wider puparium (3.89 ± 0.32 mm, range: 1.89–5.50 mm; *P* < 0.001) than *P. bennetti* did (3.50 ± 0.39 mm, range: 1.84–4.41 mm). The proportion of variance in length and width explained by both the random (*i.e*., *Farm ID* and *year*) and fixed (*i.e*., *species*) effects of the linear mixed models were 36.45% and 30.57% for the maximum puparium length and width, respectively. *Farm ID* and *year* alone accounted for 51.58% and 44.16% of these proportions, respectively.

**Figure 4.**
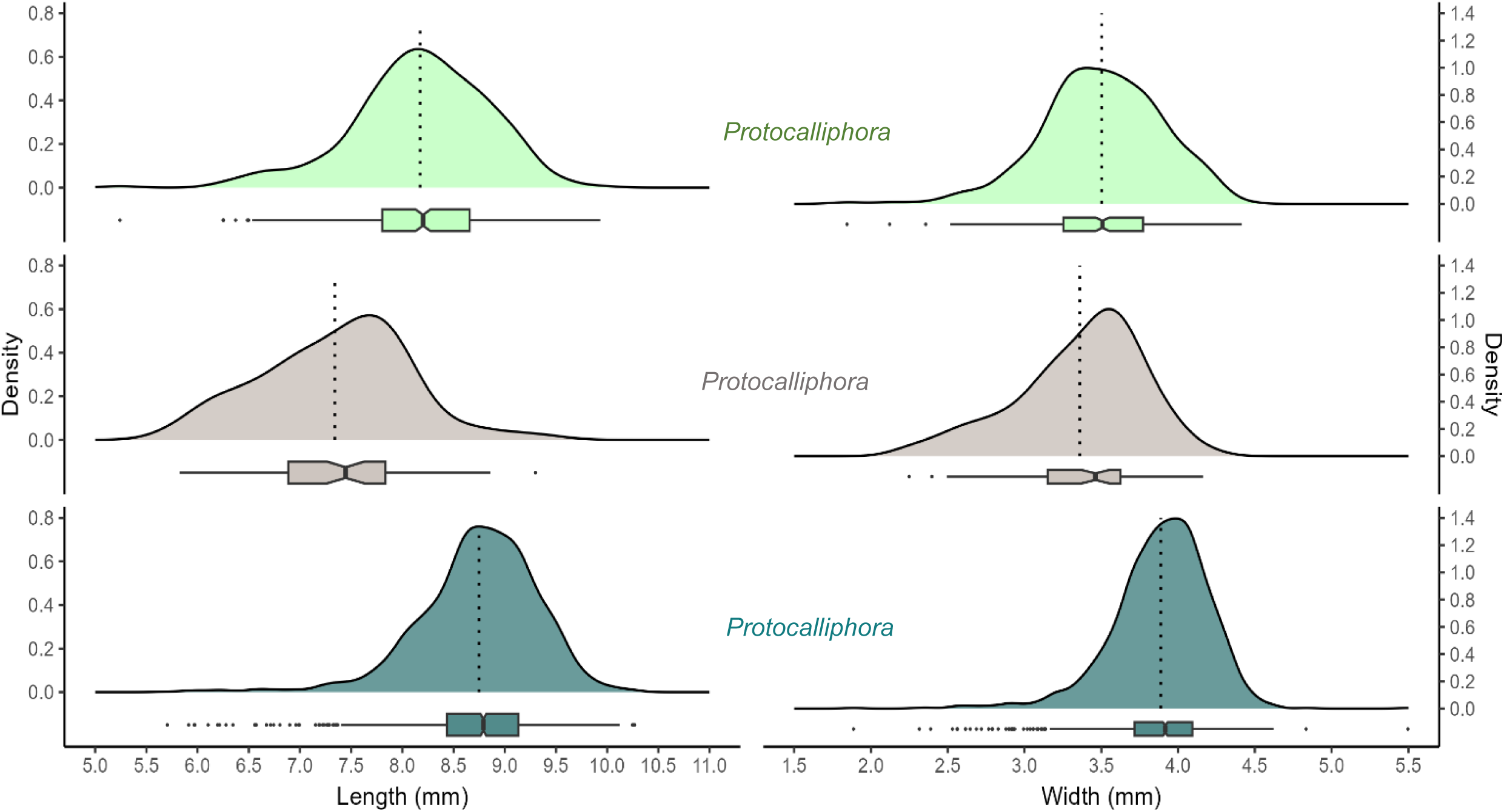
Density distribution representing the morphometric measurements (length and width in millimetres) of puparia for *Proocalliphora bennetti* (light green, n = 347), *P. metallica* (grey, n = 66), and *P. sialia* (dark green, n = 1,569) collected in southern Quebec, Canada, between 2004 and 2019 (excluding 2008–2009). Mean (dotted line) and quartiles (*via* boxplot) are shown.

## Discussion

Our study presents a rare time series on the prevalence of bird nest infestation by bird blowflies and their abundance within infested nests. We found that infestation rates and loads of *Protocalliphora* spp. within tree swallow nests were high over a 16-year period (2004–2019), yet highly variable in both space and time along a 10 200-km^2^ gradient of agricultural intensity in southern Quebec, Canada. Moreover, we observed that these fluctuations potentially followed a recurrent pattern that was shared by approximately 75% of the sampled locations (*i.e.*, farms), suggesting that both large-scale and local conditions or processes may be at play. Our study also shows that, although bird blowfly pupae were subjected to parasitoidic pressures by *Nasonia* spp. wasps that can vary as much as double among years, these were high enough to kill more than half of the *Protocalliphora* spp. pupae produced in most years. Such an impact suggests that *Protocalliphora* spp. and *Nasonia* spp. may be part of host–parasitoid dynamics that are responsible for the oscillations in bird blowfly infestation rates and loads that we report. Consistent with the previous 2-year investigation in our study system by Daoust *et al*. (2012a, 2012b), we found both mono- and multispecific infestations in nests, involving three species of *Protocalliphora* spp., with *P. sialia* dominating (> 80% of puparia). However, we found higher proportions of *P. bennetti* (∼2×) and *P. metallica* (∼4×) than reported by Daoust *et al*. (2012a), emphasising the value of multiyear studies in revealing the complete range of species composition variability in these flies. Lastly, we report highly variable puparium length and width with overlapping frequency distributions among *Protocalliphora* species. This finding suggests that these morphometric traits are not reliable as discrete diagnostic characters and should not to be used in isolation for identifying *Protocalliphora* species (Whitworth 2002). It also raises important questions about the ecological factors driving this variability and the potential consequences of size variation for emergence success and host selection by parasitoids.

### Spatiotemporal variation in *Protocalliphora* spp. prevalence and load

The prevalence and intensity of bird nest infestations by *Protocalliphora* spp. have generally been found to vary widely in both space and time. For example, Bennett and Whitworth’s (1992) comprehensive examination across 73 bird species and 4668 sampled nests revealed infestation rates of 52% within the Rocky Mountain region (Utah, United States of America) *versus* 24% in the Great Lakes region (Ontario, Canada). The mechanisms underlying such variation likely involve multiple dependent and interactive factors, and few studies have attempted to quantify the importance and spatiotemporal scales at which they operate. Although large-scale geographic variation may result from differences in climate, landscape habitat composition, bird host community (species composition and abundance), and parasitoid pressure (Sabrosky *et al*. 1989; Bennett and Whitworth 1992; Daoust *et al*. 2012a, 2012b; Garrido-Bautista *et al*. 2020), local variation may originate in factors as specific as bird host brood size, bird host hatching date (Williams 2017; Wolfe-Merritt *et al*. 2022), and nest type and material affecting the nest microclimate (Dawson *et al*. 2011; Griebel *et al*. 2020). With respect to tree swallows breeding within nest boxes, previous research reported *Protocalliphora* spp. infestation rates ranging from 89% to 100% in most study sites across Canada (Rendell and Verbeek 1996; Dawson *et al*. 2005; Gentes *et al*. 2007; Griebel *et al*. 2020). Those reported infestation rates are generally much higher than those we found either across years or farms within our study area (overall rate = 45.90%). In contrast to other study sites, many of which were located near wetlands, ours were located on farms distributed along a gradient of agricultural intensity with little, if any, wetland cover (Garrett *et al*. 2022a). Because of this, we hypothesise that the much lower infestation rates we observed are linked to intensive agricultural practices that may impact *Protocalliphora* spp*. via* multiple pathways, including directly through wetland drainage or pesticide use or indirectly by reducing the density of floral resources and bird hosts (*e.g*., Robert *et al*. 2019; Wagner *et al*. 2021; Powell *et al*. 2024). Evidence for such negative impacts has also been reported in our study area by Daoust *et al*. (2012b), who found lower loads of *Protocalliphora* spp. in tree swallow nests located within more agro-intensively worked landscapes during a 2-year period (2008–2009).

Annual infestation rates of tree swallow nests by *Protocalliphora* spp. varied between 29% and 63% across the 16 years of our study. This level of interannual variation, which was observed across the entire study area (*i.e*., 10 200 km^2^), was comparable to what we observed among nest boxes from a single farm (*i.e*., < 1 km^2^; Supplementary material, File S5). Wesołowski (2001) also reported highly variable annual nest infestation rates by *Protocalliphora* spp. in marsh tits, *Parus palustris* (Linnaeus) (Passeriformes: Paridae), that ranged from 27% to 88% in four 0.5-km^2^ plots separated by 1–2 km during an 8-year period in eastern Poland. Similarly, Musgrave *et al*. (2019) found annual infestation rates between 30% and 91% in western bluebird, *Sialia mexicana* Swainson (Passeriformes: Turdidae), nests monitored over 17 years in a an approximately 100-km^2^ area of northern New Mexico, United States of America. Despite potential confounding effects, the above comparisons suggest that nest infestation rates by *Protocalliphora* spp. typically vary substantially among years across a range of spatial scales. Given the lack of reported time series of infestation rates and loads in the literature (but see Merino *et al*. 2025), it remains largely unknown how these variables are structured in time and how such temporal structures vary across spatial scales and ecological contexts, hindering investigation into the ecological mechanisms underlying their origin. Our observation that both yearly fluctuations in bird blowfly nest infestation rates and loads were to some degree recurrent and synchronous among approximately 75% of the farms sampled over 10 200 km^2^ nonetheless suggests that regional (*e.g*., climatic), local (*e.g*., crop rotations), and density-dependent (*e.g*., parasitoidic pressure by *Nasonia* spp.) processes may be at play.

The observed loads of *Protocalliphora* spp. within infested nests exhibited a right-skewed distribution, a phenomenon commonly observed in the field in parasitology (Wilson and Grenfell 1997). Furthermore, a high degree of variability between nests, between sites, and across years have been reported (Griebel and Dawson 2019; Remeš and Krist 2005). Part of the interannual variation is often associated with weather (Merino and Potti 1996; Dawson *et al*. 2005; Castaño-Vazquez *et al*. 2021; Mennerat *et al*. 2021; Merino *et al*. 2025), and spatial variation is typically linked to micro-habitat and nest-specific factors, such as the amount of feather lining or the species and number of nestling hosts (Eeva *et al*. 1994; Tomás *et al*. 2007; Dawson *et al*. 2011; Williams 2017; Wolfe-Merritt *et al*. 2022). In our study system, part of the between-farm variation likely results from the diversified agricultural contexts in which tree swallows breed. Daoust *et al*. (2012b) found fewer *Protocalliphora* spp. in tree swallow nests located in more agro-intensive areas, particularly in maize and soybean monocultures. This pattern may result from several direct and indirect effects associated with intensive agriculture, such as reduced shelter and food resources for adult flies, lower avian host availability at both the landscape and brood scale, and increased toxicological effects from pesticide exposure either directly or through their avian hosts (Daoust *et al*. 2012b). Because *Protocalliphora* spp. loads are typically inferred from puparia counts after nestlings have left their nests (Jánošková *et al*. 2010), estimates reflect only the fraction of individuals that successfully pupated. In our case, they may underestimate the prevalence and load of *Protocalliphora* spp. in harsher environments. Given the scarcity of studies that have examined the effects of agricultural intensity on ectoparasites of wild birds, more extensive and long-term research is needed to determine the mechanisms by which agriculture generates spatiotemporal variation and ultimately shapes host–parasite dynamics.

### Parasitoidism of *Protocalliphora* spp. by *Nasonia* spp

Interannual variation in the proportions of puparia infested by *Nasonia* spp. showed no clear patterns while ranging between 48.25% and 90.48% among years, with an overall rate of 68.82%. These proportions are generally higher from those reported by Jamriška and Lučeničová (2019), who observed an average parasitoidism rate of 37.91% over four years in Slovakia, and those reported by Bennett and Whitworth (1991), who reported an overall rate of about 17% from Utah and Ontario, in North America. In contrast, Grillenberger *et al*. (2009) reported a rate of approximately 44% in New York State, United States of America, and Garrido-Bautista *et al*. (2020) reported rates from 47% to 56% for southern Spain and from 61% to 79% for central Spain, figures much closer to our own. Although more extensive longitudinal and ecological studies such as González-Bernardo *et al*. (2025) are needed, current findings suggest that geographical and large-scale environmental contexts may significantly influence the likelihood of *Nasonia* spp. parasitoidism (Garrido-Bautista *et al*. 2024), a primary determinant of *Protocalliphora* spp. emergence success. However, it is crucial to recognise that, in addition to the substantial control exerted by *Nasonia* spp. on bird blowflies communities, uninfested *Protocalliphora* spp. individuals may also die during pupation due to other ecological factors, such as environmental contamination. The specific impacts of those factors on *Protocalliphora* spp. emergence success have yet to be thoroughly investigated and quantified.

*Nasonia* spp. parasitoid wasps have mostly served as model species in numerous laboratory studies addressing themes in genetics, developmental biology, and behavioural ecology (Werren and Loehlin 2009; Niehuis *et al*. 2010; Nadal-Jimenez *et al*. 2023). The ecological factors driving host selection and parasitoidism by *Nasonia* spp. under natural conditions are thus poorly understood (Peters 2010; Peters and Abraham 2010; Garrido-Bautista *et al*. 2020, 2024). This is particularly true given the wide diversity of hosts in which *Nasonia* spp. can lay eggs and the potential for regional variations due to their widespread distribution (Skovgård and Jespersen 1999; Oliva 2008). This leaves the causes of yearly fluctuations in *Protocalliphora* spp. puparia parasitoidic events and of winter diapause within the host puparia unresolved. Future research on *Nasonia* spp. ecology would likely benefit from considering factors such as vegetation type and density, because these features may influence host parasitoidism by affecting the availability of food and shelter (Wylie 1958). Avian nest availability associated with these environmental factors could also potentially induce host-dependent effects, contributing to an increased incidence of *Nasonia* spp. parasitoidism on *Protocalliphora* spp.

The present study reports the first quantitative exploration of dormancy and mortality of *Nasonia* spp. within their host in the wild. We noted consistently low yearly proportions of *Protocalliphora* spp. puparia containing dormant or dead *Nasonia* spp. (range: 3.07–16.36%), with no discernible temporal trend. Several biotic and abiotic factors, such as weather, luminosity, or host and parasitoid body condition, may affect the production of diapausing *Nasonia* spp. offspring (Saunders 1969; Yoder *et al*. 1994; Bertossa *et al*. 2010; Paolucci *et al*. 2013; Li *et al*. 2014). Mechanisms linked to phenology could thereby affect the dormancy and survival of *Nasonia* spp. larvae. For instance, bird blowfly pupation phenology, which in turn determines host availability and development conditions for *Nasonia* spp., must partly depend on avian host breeding phenology. Moreover, agricultural intensity may affect *Nasonia* spp. development and emergence success because pesticide use can alter parasitoid wasps’ cognition and fitness (Schöfer *et al*. 2023, 2024; Sulg *et al*. 2023) and the availability and quality of their hosts (Daoust *et al*. 2012b; Li *et al*. 2014).

### *Protocalliphora* species composition and morphometry

Bird nestlings from the Nearctic region are hosts to at least 28 *Protocalliphora* species (Sabrosky *et al*. 1989; updated by Whitworth 2002). Although some bird blowfly species appear to specialise on specific hosts through behavioural adaptations, some parasitise a wide array of bird species (Bennett and Whitworth 1992), and the occurrence of multiple *Protocalliphora* species within a single host nest is not rare (Daoust *et al*. 2012a). Our longitudinal study, one of the few that describe *Protocalliphora* species composition, reports that such compositions can vary substantially across years. Indeed, our results show that certain species (in this case, *P. metallica*) may be practically absent in some years, only to reappear in relatively high numbers the following year. Even the highly prevalent *P. sialia* fluctuated widely, representing 58.93% of the *Protocalliphora* specimens in 2011 but being almost the only species found in infested nests the following year (98.32%). The results show that longitudinal studies considering variability in species composition are essential before conclusions about host specialisation to avian parasites and mixed infestations can be drawn.

Although descriptive statistics of some morphometric measurements of *Protocalliphora* spp. puparia can be found in the literature, few, if any, studies report the actual frequency distributions of these features that may help discriminating among species of this cryptic group (Heeb *et al*. 2000; Whitworth 2002, 2003; Jamriška and Modrý 2013). The mean, standard deviation, and range estimates of puparium length and width for *P. bennetti, P. metallica,* and *P. sialia* that we obtained *via* photogrammetry align well with published estimates (Sabrosky *et al*. 1989; Whitworth 2002). However, the fact that density plots highlighted substantial intraspecific variation in both puparium length and width, as well as significant overlap among the three studied species, indicates that these metrics should not be used on their own for *Protocalliphora* spp. identification, even within a small regional species pool. Nonetheless, the ecological factors contributing to this high intraspecific variation, with certain puparia being more than twice as large and long as others, is unclear. Sexual dimorphism (Sabrosky *et al*. 1989) or factors like host availability and quality, as well as developmental period and temperature, which are often interconnected, have been hypothesised as causes of size variation in *Protocalliphora* spp. For instance, underfed larvae may conclude their development as ‘runts’ (Sabrosky *et al*. 1989; Bennett and Whitworth 1991). Alternatively, intensive agricultural practices, including pesticide use, have been reported as either direct or indirect drivers of phenotypic variation (Wisniewska and Prokopy 1997; Verheyen and Stoks 2019; Gérard *et al*. 2022; Ponce-Méndez *et al*. 2022) and pupation success (Heneberg *et al*. 2020; Mahdjoub *et al*. 2020) in nontargeted arthropod species.

All the ecological parameters investigated in the present study showed substantial variation in space and time or among individuals, indicating that future studies addressing the mechanisms driving *Protocalliphora* spp. host–parasite dynamics adopt a long-term monitoring and multitrophic approach in contrasting environments. Investigating the tri-trophic relationships involving birds, *Protocalliphora* spp., and *Nasonia* spp. across diverse agroecosystems presents a promising avenue for such research. Given the complex interplay among environmental conditions and multitrophic interactions, we suggest further that such endeavours be tackled using a causal inference framework to limit biases, especially because each trophic level may respond to different factors at different spatiotemporal scales and potentially nonlinearly or in different directions, which has the potential to amplify, attenuate, mask, and even create effects (Arif and MacNeil 2023; Grace 2024).

## Acknowledgements

The authors are deeply grateful to the many farm owners who have collaborated to our long-term monitoring program. They also thank the Université de Sherbrooke for its institutional support and the funding agencies that made this research possible, including the Natural Sciences and Engineering Research Council of Canada (NSERC), the Fonds de recherche du Québec – Nature et technologies (FRQ-NT), and the Canada Research Chairs program. They warmly acknowledge the contribution of the many students, field assistants, and technicians, and especially of L. Glaude and Y. Sageau, who participated to the collection and sorting of nests over the years. Special thanks are due D. Garneau for his generous assistance with all aspects of imagery. This study was conducted with the approval of the Université de Sherbrooke animal care committee, and the authors gratefully recognise all those whose efforts, on the ground and behind the scenes, contributed to making the work possible.

