## Supplementary Material 1 for "Parasitism by, species composition, morphometry, and parasitoidism of *Protocalliphora* bird blowflies (Diptera: Calliphoridae) in Quebec’s farmlands: a 16-year dataset"

**Supplementary Material 1 – Study area**

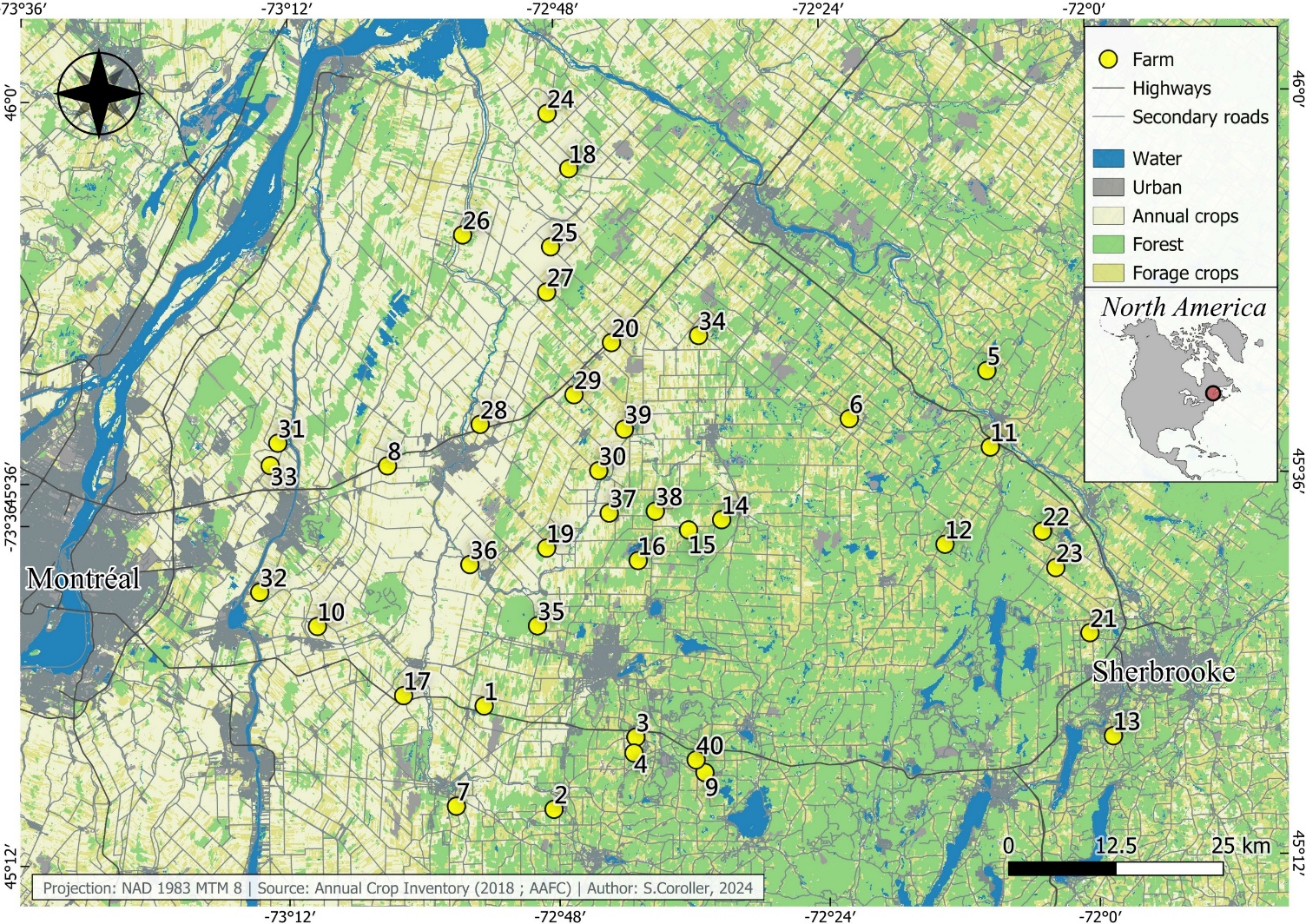
As mentioned in the section Study system and field sampling, we collected *Protocalliphora* spp. puparia within a network of 400 nest boxes distributed equally among 40 farms located along a 10,200-km² gradient of agricultural intensity in southern Québec. This area spans a vast region composed of farmlands that vary strikingly in both their landscape composition and configuration due to agricultural intensity, as can be seen in the following figure.

**Figure S1**. Geographic distribution of the 40 farms monitored across a 10,200-km² gradient of agricultural intensity in southern Québec, Canada. The reported coordinates are in the Universal Transverse Mercator (UTM) zone 18 north, while the mapping was conducted using the NAD83 MTM zone 8 system.

### **Supplementary Material 2 – Agricultural intensification and random sampling of *Protocalliphora*-infested nest boxes**

We quantified the landscape habitat composition within a 500-m radius surrounding each Tree Swallow (*Tachycineta bicolor*) nest box by combining in-situ habitat characterization with high-resolution orthophotos (1:40,000; QGIS, 2020). Habitat mapping was conducted at the end of each breeding season between 2006 and 2019 and included four major land-cover categories: Forest, Corn/Soybean, Other Cereals (wheat, oat, barley), and Pasture/Forage (hay, alfalfa, clover). Orthophotos were used to delineate field and forest boundaries, following the procedures outlined in Garrett et al. (2022a) and Powell et al. (2024). Streams and water bodies covered on average less than 1% of the area surrounding nest boxes (Garrett et al., 2022b) and were thus excluded from analyses.

To characterize the average landscape habitat composition of each farm, we performed a robust compositional Principal Component Analysis (rPCA) using the *robCompositions* R package (Templ et al., 2011) on the annual relative cover of the four habitat types within each 500-m buffer (2006–2019). The first component (Comp.1) explained 79% of the total variance and was positively correlated with Corn/Soybean and negatively with Forage and Forest cover, representing a gradient of agricultural intensity. The second component (Comp.2) explained 16% of the variance and was positively associated with Tree cover and negatively with Forage, representing a tree–forage gradient. As in-situ characterization of crops could not be performed in 2004 and 2005, Comp.1 and Comp.2 values for those years were estimated by averaging the corresponding 2006–2008 scores for each nest box. This approach was justified by the relatively stable landscape configuration observed over time within farms, although interannual changes in field boundaries and crop rotations were not negligible and can contribute meaningfully to local habitat heterogeneity.

To select a representative subset of nests infested by *Protocalliphora* spp. for further analyses of their puparia, we first stratified the full dataset according to the 33% and 66% quantiles of both Comp.1 and Comp.2 scores, thereby defining nine distinct classes of “agricultural context” (e.g., *Low Comp.1 × Low Comp.2*, *Mid Comp.1 × High Comp.2*, etc.; Fig. S2.A). One farm was then randomly selected within each of these nine agricultural-context classes before an additional farm was randomly chosen among the remaining ones, for a total of 10 farms. Because each farm contained 10 nest boxes spaced by ~50 m, the landscape habitat composition surrounding each box varied slightly within farms. Moreover, as field configuration and crop composition could shift from year to year due to crop rotations and local management practices, randomly selecting two infested nests (when available) per farm allowed for substantial landscape heterogeneity across the time series (Fig. S2B). To ensure adequate representation across years and maintain statistical robustness, we sometimes had to randomly select additional farms. We randomly selected up to two *Protocalliphora*-infested nests from these additional farms and stopped as soon as we could process ≥ 100 *Protocalliphora* spp. puparia on each year. A total of 263 nests were thereby subsampled across farms and years, representing a broad spectrum of agricultural intensity and landscape structural complexity among the 1,227 nests found to be infested by *Protocalliphora* spp. between 2004 and 2019 in our study system.

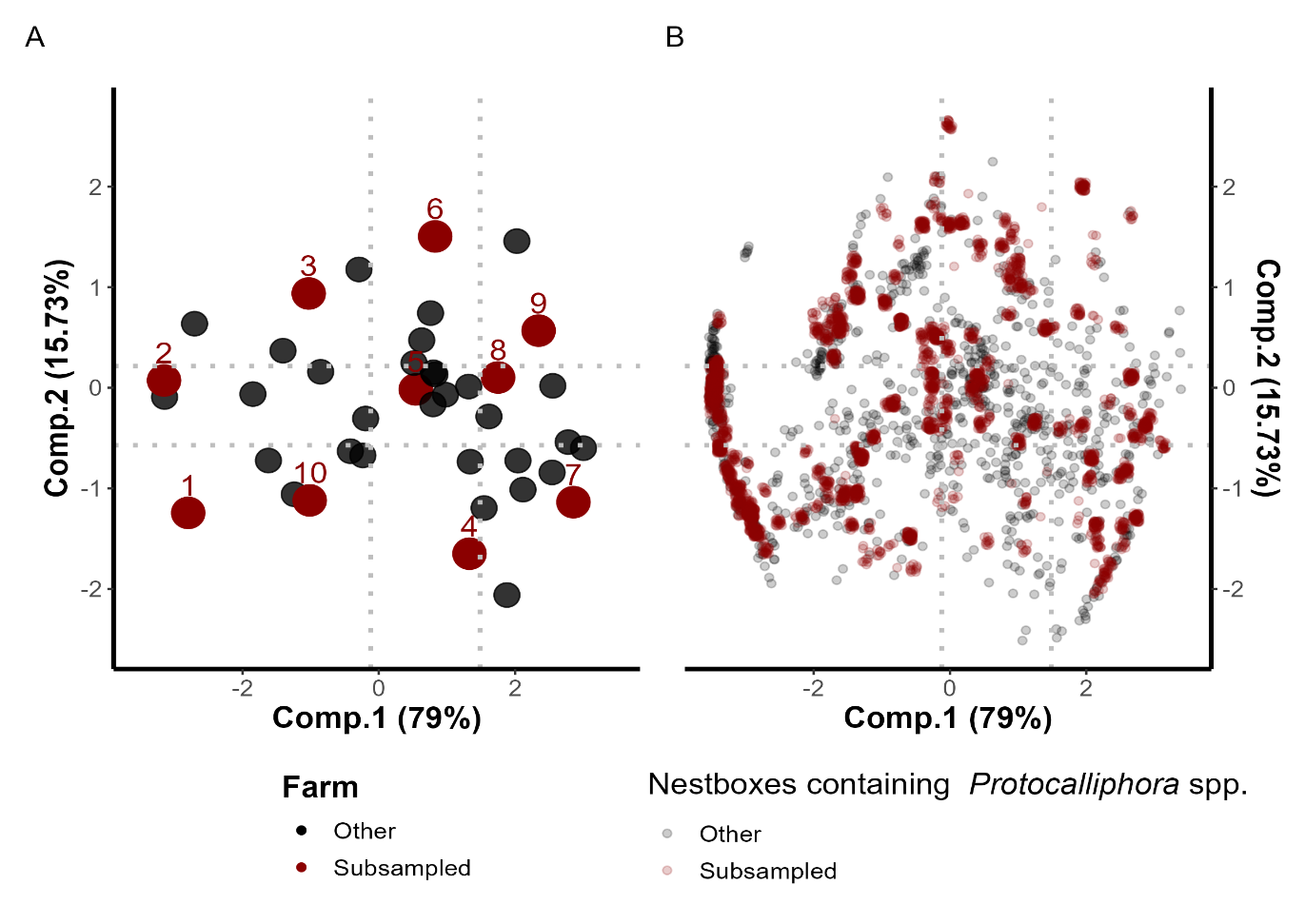
**Figure S2.** (A) Stratification of the 40 farms across the agricultural landscape gradient defined by a robust compositional PCA (rPCA) of habitat types within 500 m of nest boxes. Grey dashed lines mark the 33% and 66% quantiles of Comp.1 and Comp.2 scores, defining nine classes of agricultural context. Among the 40 farms (black dots), 10 (red dots) were randomly selected across these classes, with up to two *Protocalliphora*-infested nests per farm and year. Additional randomly selected farms and nests (≤ 2 nests) were sequentially added to ensure that ≥ 100 puparia could be processed on each year, increasing coverage of landscape contexts. (B) Distribution of the 263 nests (red dots) selected among all 1,227 nests infested by *Protocalliphora* spp. (grey dots) between 2004 and 2019.

### **Supplementary Material 3 – Model diagnostics of temporal trends in *Protocalliphora* spp. nest infestation and loads**

This second Supplementary Material reports the codes, model diagnostics and results of the Generalized Linear Additive Mixed Models (GAMMs) fitted with the *mgcv* package (v.1.9-1; Wood 2023) in R (v.4.3.1, R Core Team, 2023) and used to explore temporal trends in the infestation rate of Tree Swallow nests by *Protocalliphora* and the number of *Protocalliphora* puparia per infested nest found on the 40 farms that were monitored between 2004 and 2019. We assessed how the annual mean nest infestation rate across farms varied during this period, and to what level each farm followed that global trend, if any. We modeled the response variable using a global smooth term for Year and group-level smooth terms allowing for farm-specific temporal trends that may differ in shape and wiggliness from the global trend (i.e., see Model *GI* in Pedersen et al., 2019). Temporal autocorrelation in the residuals was assessed using the *acf* function in R (Simpson, 2018).

**Script S3**. Codes, model diagnostics and results of the Generalized Linear Additive Mixed Models (GAMMs) fitted with the *mgcv* package (v.1.9-1; Wood 2023) in R (v.4.3.1, R Core Team, 2023.

**Nest infestation rate**

```
m_rate <- gam(Infest.rate ~ s(Year, k=15, bs="tp", m=2) + s(Year, by=Farm, k=10, bs="tp", m=1) + s(Farm, bs="re", k=20), family=binomial(link="logit"), weight=n_Broods, data=data.rate, method="ML")

summary(m_rate)
##
## Family: binomial
## Link function: logit
##
## Formula:
## Infest.rate ~ s(Year, k = 15, bs = "tp", m = 2) + s(Year, by = Farm,
##     k = 10, bs = "tp", m = 1) + s(Farm, bs = "re", k = 20)
##
## Parametric coefficients:
##             Estimate Std. Error z value Pr(>|z|)
## (Intercept)  -0.2834     0.1376   -2.06   0.0394 *
## ---
## Signif. codes:  0 '***' 0.001 '**' 0.01 '*' 0.05 '.' 0.1 ' ' 1
##
## Approximate significance of smooth terms:
##                      edf Ref.df  Chi.sq p-value
## s(Year)        1.298e+01  13.81 101.401 < 2e-16 ***
## s(Year):Farm1  1.432e+00   9.00   2.988 0.11949
## s(Year):Farm2  3.708e+00   9.00  22.533 0.12997
## s(Year):Farm3  6.559e+00   9.00  42.325 0.00259 **
## s(Year):Farm4  7.035e-05   9.00   0.000 0.61305
## s(Year):Farm5  1.433e-05   9.00   0.000 0.68286
## s(Year):Farm6  4.148e-05   9.00   0.000 0.91376
## s(Year):Farm7  2.307e-04   9.00   0.000 0.38550
## s(Year):Farm8  4.943e-06   8.00   0.000 0.42310
## s(Year):Farm9  6.290e-05   9.00   0.000 0.50425
## s(Year):Farm10 1.732e+00   9.00   3.169 0.17457
## s(Year):Farm11 5.248e-05   9.00   0.000 0.69822
## s(Year):Farm12 4.475e-01   9.00   0.564 0.25871
## s(Year):Farm13 2.932e-01   9.00   0.342 0.30346
## s(Year):Farm14 1.848e-04   9.00   0.000 0.48403
## s(Year):Farm15 2.432e+00   9.00   6.601 0.03064 *
## s(Year):Farm16 8.780e-05   9.00   0.000 0.58275
## s(Year):Farm17 2.734e+00   9.00  11.546 0.00378 **
## s(Year):Farm18 2.608e-05   9.00   0.000 0.80270
## s(Year):Farm19 4.590e-05   9.00   0.000 0.65308
## s(Year):Farm20 4.753e-04   9.00   0.000 0.38212
## s(Year):Farm21 3.433e-05   9.00   0.000 0.90616
## s(Year):Farm22 2.879e+00   9.00  19.168 0.00530 **
## s(Year):Farm23 6.771e-05   9.00   0.000 0.89045
## s(Year):Farm24 2.125e+00   9.00   4.622 0.05964 .
## s(Year):Farm25 2.857e-05   9.00   0.000 0.93904
## s(Year):Farm26 1.623e-04   9.00   0.000 0.46858
## s(Year):Farm27 8.269e-05   9.00   0.000 0.74348
## s(Year):Farm28 4.446e-05   9.00   0.000 0.73338
## s(Year):Farm29 4.268e-01   9.00   0.637 0.23083
## s(Year):Farm30 2.055e+00   9.00   3.952 0.24409
## s(Year):Farm31 5.145e-05   9.00   0.000 0.69437
## s(Year):Farm32 3.899e+00   9.00  15.467 0.11586
## s(Year):Farm33 5.940e+00   9.00  14.609 0.02290 *
## s(Year):Farm34 1.612e+00   9.00   4.686 0.03724 *
## s(Year):Farm35 1.246e-05   9.00   0.000 0.83202
## s(Year):Farm36 2.341e+00   9.00   6.447 0.05096 .
## s(Year):Farm37 1.169e+00   9.00   2.344 0.09712 .
## s(Year):Farm38 5.650e+00   9.00  14.580 0.01924 *
## s(Year):Farm39 3.222e+00   9.00  14.189 0.01506 *
## s(Year):Farm40 1.645e+00   9.00   4.159 0.04680 *
## s(Farm)        3.336e+01  39.00 254.388 < 2e-16 ***
## ---
## Signif. codes:  0 '***' 0.001 '**' 0.01 '*' 0.05 '.' 0.1 ' ' 1
##
## R-sq.(adj) =  0.488   Deviance explained = 49.8%
## -ML =  864.2  Scale est. = 1         n = 587
```

gam.check(m_rate)

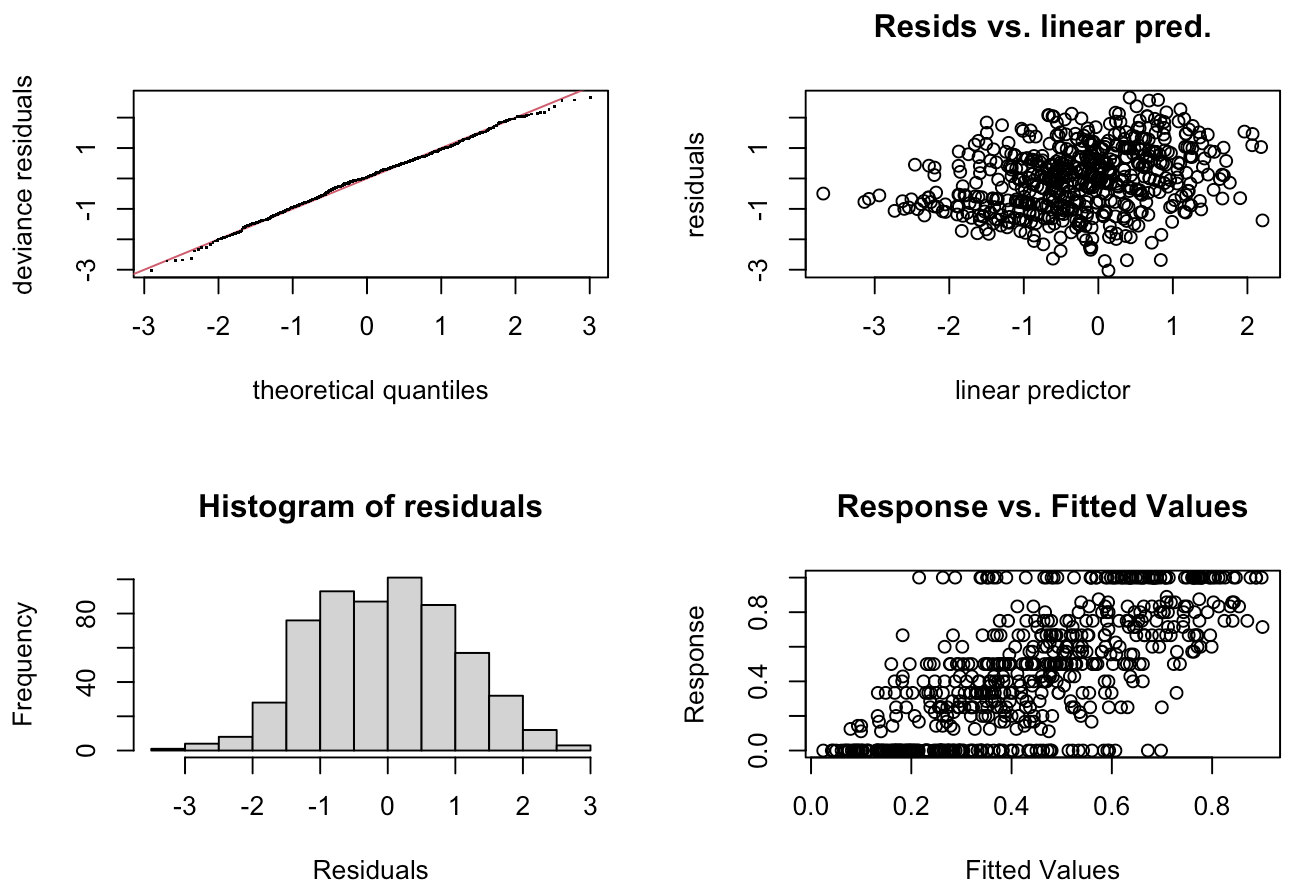

```
##
## Method: ML   Optimizer: outer newton
## full convergence after 13 iterations.
## Gradient range [-2.552142e-05,2.841186e-05]
## (score 864.196 & scale 1).
## Hessian positive definite, eigenvalue range [4.343004e-07,13.99244].
## Model rank =  415 / 415
##
## Basis dimension (k) checking results. Low p-value (k-index<1) may
## indicate that k is too low, especially if edf is close to k'.
##
##                      k'      edf k-index p-value
## s(Year)        1.40e+01 1.30e+01    0.99    0.46
## s(Year):Farm1  9.00e+00 1.43e+00    0.99    0.45
## s(Year):Farm2  9.00e+00 3.71e+00    0.99    0.43
## s(Year):Farm3  9.00e+00 6.56e+00    0.99    0.36
## s(Year):Farm4  9.00e+00 7.04e-05    0.99    0.39
## s(Year):Farm5  9.00e+00 1.43e-05    0.99    0.38
## s(Year):Farm6  9.00e+00 4.15e-05    0.99    0.36
## s(Year):Farm7  9.00e+00 2.31e-04    0.99    0.43
## s(Year):Farm8  9.00e+00 4.94e-06    0.99    0.43
## s(Year):Farm9  9.00e+00 6.29e-05    0.99    0.39
## s(Year):Farm10 9.00e+00 1.73e+00    0.99    0.46
## s(Year):Farm11 9.00e+00 5.25e-05    0.99    0.46
## s(Year):Farm12 9.00e+00 4.48e-01    0.99    0.40
## s(Year):Farm13 9.00e+00 2.93e-01    0.99    0.42
## s(Year):Farm14 9.00e+00 1.85e-04    0.99    0.50
## s(Year):Farm15 9.00e+00 2.43e+00    0.99    0.46
## s(Year):Farm16 9.00e+00 8.78e-05    0.99    0.43
## s(Year):Farm17 9.00e+00 2.73e+00    0.99    0.47
## s(Year):Farm18 9.00e+00 2.61e-05    0.99    0.42
## s(Year):Farm19 9.00e+00 4.59e-05    0.99    0.41
## s(Year):Farm20 9.00e+00 4.75e-04    0.99    0.48
## s(Year):Farm21 9.00e+00 3.43e-05    0.99    0.43
## s(Year):Farm22 9.00e+00 2.88e+00    0.99    0.43
## s(Year):Farm23 9.00e+00 6.77e-05    0.99    0.45
## s(Year):Farm24 9.00e+00 2.12e+00    0.99    0.42
## s(Year):Farm25 9.00e+00 2.86e-05    0.99    0.39
## s(Year):Farm26 9.00e+00 1.62e-04    0.99    0.40
## s(Year):Farm27 9.00e+00 8.27e-05    0.99    0.44
## s(Year):Farm28 9.00e+00 4.45e-05    0.99    0.42
## s(Year):Farm29 9.00e+00 4.27e-01    0.99    0.44
## s(Year):Farm30 9.00e+00 2.05e+00    0.99    0.34
## s(Year):Farm31 9.00e+00 5.14e-05    0.99    0.35
## s(Year):Farm32 9.00e+00 3.90e+00    0.99    0.43
## s(Year):Farm33 9.00e+00 5.94e+00    0.99    0.40
## s(Year):Farm34 9.00e+00 1.61e+00    0.99    0.40
## s(Year):Farm35 9.00e+00 1.25e-05    0.99    0.42
## s(Year):Farm36 9.00e+00 2.34e+00    0.99    0.39
## s(Year):Farm37 9.00e+00 1.17e+00    0.99    0.42
## s(Year):Farm38 9.00e+00 5.65e+00    0.99    0.45
## s(Year):Farm39 9.00e+00 3.22e+00    0.99    0.40
## s(Year):Farm40 9.00e+00 1.65e+00    0.99    0.44
## s(Farm)        4.00e+01 3.34e+01      NA      NA


acf(residuals(m_rate), lag.max = 10, main = "ACF")
```

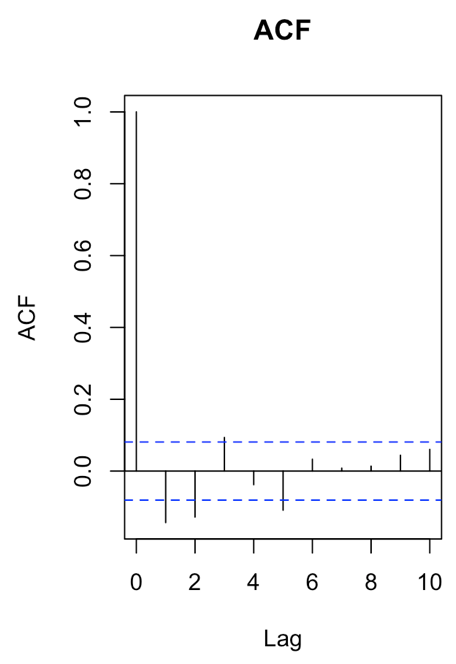

draw(m_rate, select=c(1,42))

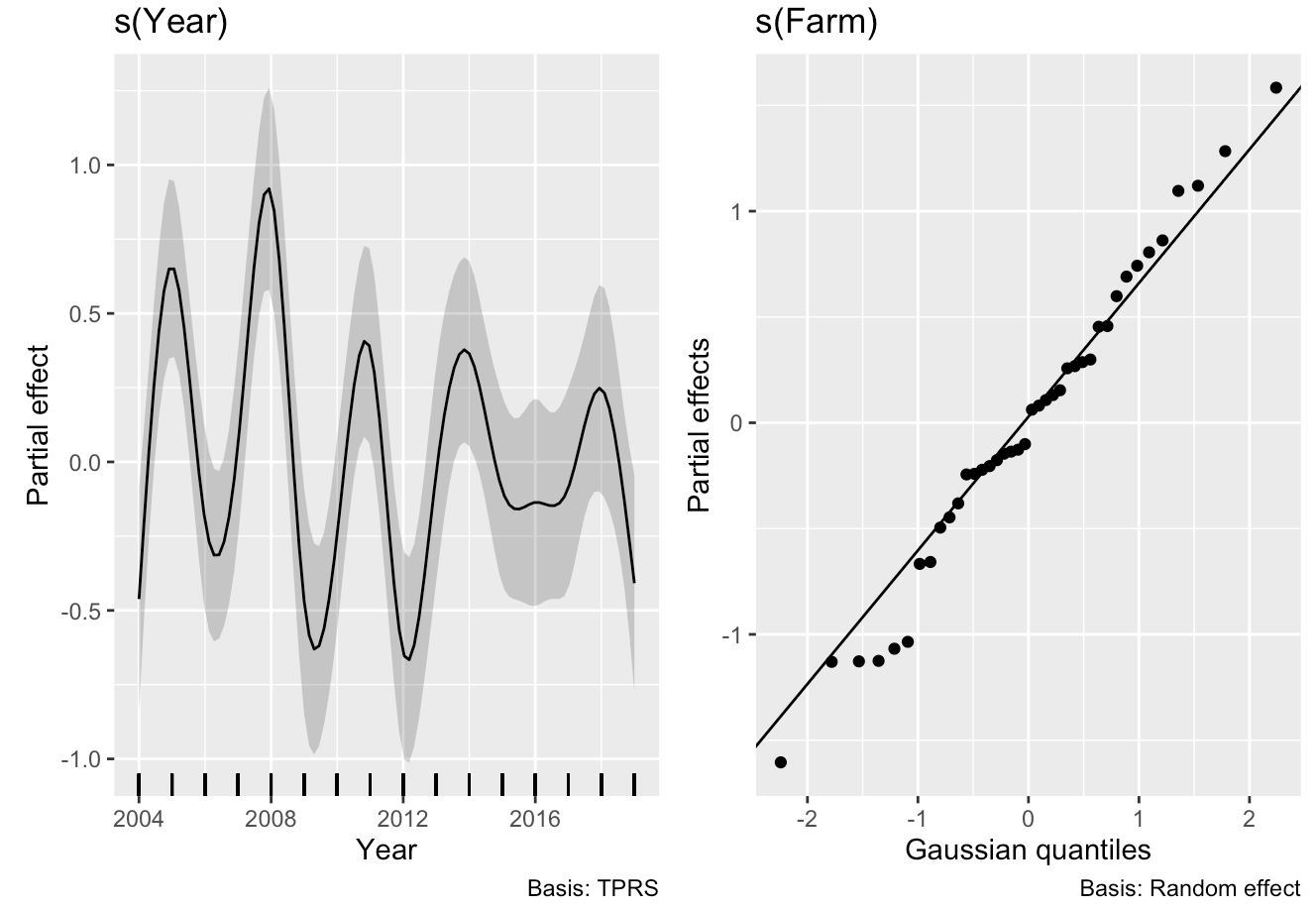

**Number of *Protocalliphora* puparia per infested nest**

```
m_number <- gam(n_Protos ~ s(Year, k=15, bs="tp", m=2) + s(Year, by=Farm, k=10, bs="tp", m=1) + s(Farm, bs="re", k=20), family=tw, data=data.load.infest, method="ML")

summary(m_number)
## Family: Tweedie(p=1.99)
## Link function: log
##
## Formula:
## n_Protos ~ s(Year, k = 15, bs = "tp", m = 2) + s(Year, by = Farm,
##     k = 10, bs = "tp", m = 1) + s(Farm, bs = "re", k = 20)
##
## Parametric coefficients:
##             Estimate Std. Error t value Pr(>|t|)
## (Intercept)  2.76387    0.04958   55.75   <2e-16 ***
## ---
## Signif. codes:  0 '***' 0.001 '**' 0.01 '*' 0.05 '.' 0.1 ' ' 1
##
## Approximate significance of smooth terms:
##                      edf Ref.df     F  p-value
## s(Year)        1.138e+01  12.92 7.012  < 2e-16 ***
## s(Year):Farm1  1.373e-02   9.00 0.001  0.62802
## s(Year):Farm2  8.747e-01   6.00 0.284  0.30627
## s(Year):Farm3  1.382e-03   9.00 0.000  0.57734
## s(Year):Farm4  7.937e-01   9.00 0.148  0.15892
## s(Year):Farm5  1.779e-03   9.00 0.000  0.47290
## s(Year):Farm6  1.095e-02   9.00 0.001  0.27087
## s(Year):Farm7  7.159e-04   9.00 0.000  0.61489
## s(Year):Farm8  9.436e-04   5.00 0.000  0.83031
## s(Year):Farm9  1.623e+00   9.00 0.592  0.01986 *
## s(Year):Farm10 2.376e-03   9.00 0.000  0.38974
## s(Year):Farm11 9.511e-01   9.00 0.214  0.10781
## s(Year):Farm12 1.286e+00   9.00 0.234  0.15730
## s(Year):Farm13 7.598e-01   9.00 0.123  0.20101
## s(Year):Farm14 9.879e-01   9.00 0.176  0.16614
## s(Year):Farm15 1.947e-03   9.00 0.000  0.59872
## s(Year):Farm16 2.497e-03   9.00 0.000  0.55083
## s(Year):Farm17 4.124e-01   9.00 0.052  0.30715
## s(Year):Farm18 8.711e-04   9.00 0.000  0.83578
## s(Year):Farm19 1.514e+00   6.00 0.763  0.03419 *
## s(Year):Farm20 3.009e+00   7.00 1.497  0.01188 *
## s(Year):Farm21 9.937e-04   9.00 0.000  0.78825
## s(Year):Farm22 2.230e+00   9.00 0.820  0.01336 *
## s(Year):Farm23 1.350e+00   9.00 0.224  0.19458
## s(Year):Farm24 1.159e-03   9.00 0.000  0.59947
## s(Year):Farm25 1.427e+00   3.00 2.255  0.00748 **
## s(Year):Farm26 4.506e-04   9.00 0.000  0.99107
## s(Year):Farm27 2.652e-03   9.00 0.000  0.66958
## s(Year):Farm28 2.360e+00   7.00 2.994 3.24e-05 ***
## s(Year):Farm29 2.537e-03   4.00 0.001  0.35355
## s(Year):Farm30 2.573e+00   8.00 0.896  0.05671 .
## s(Year):Farm31 7.689e-01   9.00 0.156  0.14029
## s(Year):Farm32 1.054e-03   9.00 0.000  0.43156
## s(Year):Farm33 8.575e-01   9.00 0.145  0.18986
## s(Year):Farm34 1.258e+00   3.00 1.558  0.04346 *
## s(Year):Farm35 1.565e-03   9.00 0.000  0.75436
## s(Year):Farm36 1.263e-02   9.00 0.001  0.46835
## s(Year):Farm37 2.482e-03   9.00 0.000  0.38497
## s(Year):Farm38 1.802e-03   9.00 0.000  0.45869
## s(Year):Farm39 1.037e+00   8.00 0.277  0.13061
## s(Year):Farm40 2.074e+00   9.00 0.761  0.01716 *
## s(Farm)        2.179e+01  39.00 1.884  < 2e-16 ***
## ---
## Signif. codes:  0 '***' 0.001 '**' 0.01 '*' 0.05 '.' 0.1 ' ' 1
##
## R-sq.(adj) =  0.122   Deviance explained = 17.2%
## -ML = 4760.3  Scale est. = 0.95101   n = 1227
```

gam.check(m_number)

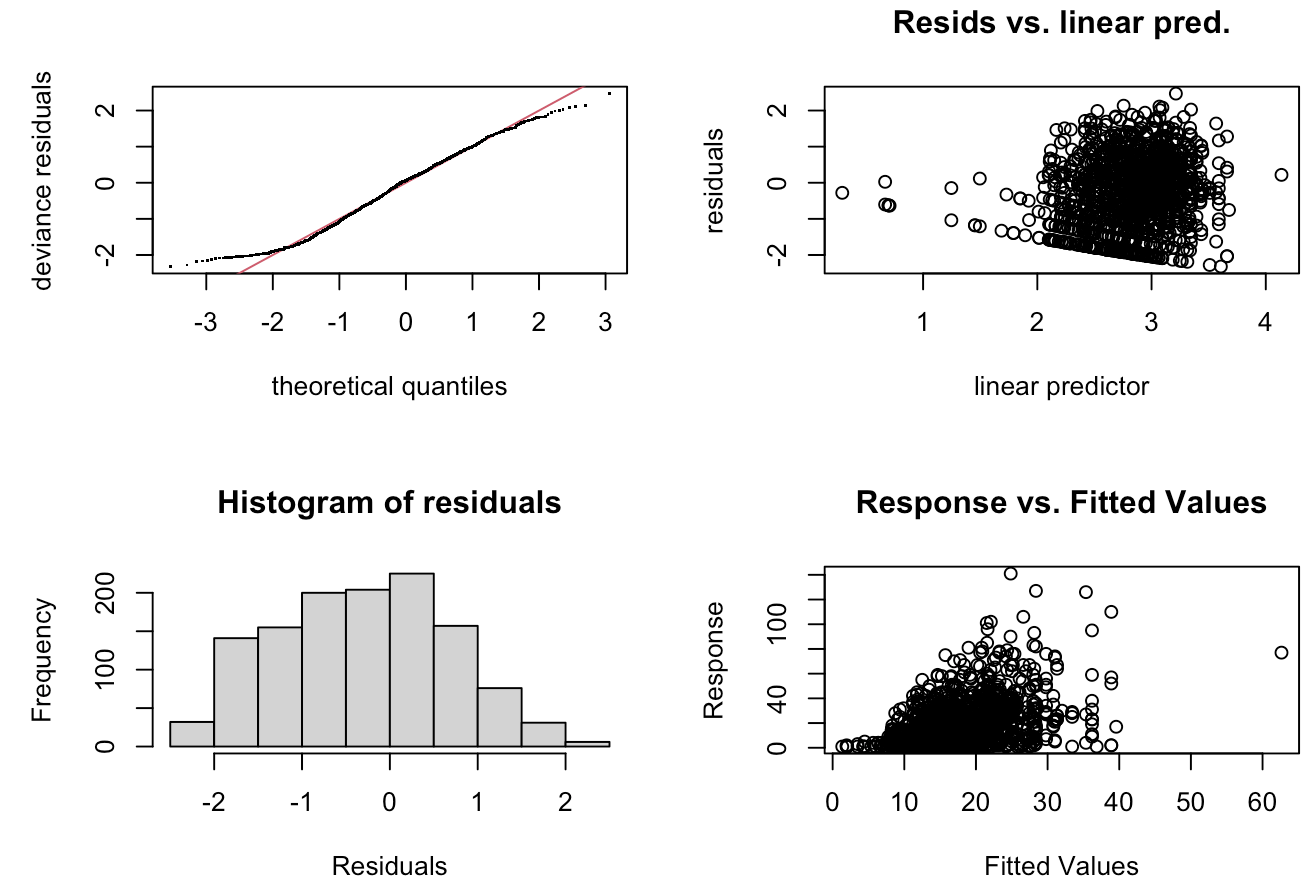

```
##
## Method: ML   Optimizer: outer newton
## full convergence after 17 iterations.
## Gradient range [-0.009328435,0.000225552]
## (score 4760.302 & scale 0.9510084).
## eigenvalue range [-0.0002268827,784.2636].
## Model rank =  415 / 415
##
## Basis dimension (k) checking results. Low p-value (k-index<1) may
## indicate that k is too low, especially if edf is close to k'.
##
##                      k'      edf k-index p-value
## s(Year)        1.40e+01 1.14e+01    0.89    0.28
## s(Year):Farm1  9.00e+00 1.37e-02    0.89    0.25
## s(Year):Farm2  9.00e+00 8.75e-01    0.89    0.25
## s(Year):Farm3  9.00e+00 1.38e-03    0.89    0.25
## s(Year):Farm4  9.00e+00 7.94e-01    0.89    0.28
## s(Year):Farm5  9.00e+00 1.78e-03    0.89    0.31
## s(Year):Farm6  9.00e+00 1.10e-02    0.89    0.30
## s(Year):Farm7  9.00e+00 7.16e-04    0.89    0.27
## s(Year):Farm8  9.00e+00 9.44e-04    0.89    0.24
## s(Year):Farm9  9.00e+00 1.62e+00    0.89    0.25
## s(Year):Farm10 9.00e+00 2.38e-03    0.89    0.21
## s(Year):Farm11 9.00e+00 9.51e-01    0.89    0.28
## s(Year):Farm12 9.00e+00 1.29e+00    0.89    0.21
## s(Year):Farm13 9.00e+00 7.60e-01    0.89    0.25
## s(Year):Farm14 9.00e+00 9.88e-01    0.89    0.28
## s(Year):Farm15 9.00e+00 1.95e-03    0.89    0.30
## s(Year):Farm16 9.00e+00 2.50e-03    0.89    0.23
## s(Year):Farm17 9.00e+00 4.12e-01    0.89    0.21
## s(Year):Farm18 9.00e+00 8.71e-04    0.89    0.28
## s(Year):Farm19 9.00e+00 1.51e+00    0.89    0.28
## s(Year):Farm20 9.00e+00 3.01e+00    0.89    0.26
## s(Year):Farm21 9.00e+00 9.94e-04    0.89    0.24
## s(Year):Farm22 9.00e+00 2.23e+00    0.89    0.20
## s(Year):Farm23 9.00e+00 1.35e+00    0.89    0.22
## s(Year):Farm24 9.00e+00 1.16e-03    0.89    0.26
## s(Year):Farm25 9.00e+00 1.43e+00    0.89    0.26
## s(Year):Farm26 9.00e+00 4.51e-04    0.89    0.27
## s(Year):Farm27 9.00e+00 2.65e-03    0.89    0.31
## s(Year):Farm28 9.00e+00 2.36e+00    0.89    0.24
## s(Year):Farm29 9.00e+00 2.54e-03    0.89    0.25
## s(Year):Farm30 9.00e+00 2.57e+00    0.89    0.28
## s(Year):Farm31 9.00e+00 7.69e-01    0.89    0.26
## s(Year):Farm32 9.00e+00 1.05e-03    0.89    0.22
## s(Year):Farm33 9.00e+00 8.57e-01    0.89    0.28
## s(Year):Farm34 9.00e+00 1.26e+00    0.89    0.28
## s(Year):Farm35 9.00e+00 1.56e-03    0.89    0.21
## s(Year):Farm36 9.00e+00 1.26e-02    0.89    0.23
## s(Year):Farm37 9.00e+00 2.48e-03    0.89    0.26
## s(Year):Farm38 9.00e+00 1.80e-03    0.89    0.28
## s(Year):Farm39 9.00e+00 1.04e+00    0.89    0.28
## s(Year):Farm40 9.00e+00 2.07e+00    0.89    0.21
## s(Farm)        4.00e+01 2.18e+01      NA      NA


acf(residuals(m4), lag.max = 10, main = "ACF")
```

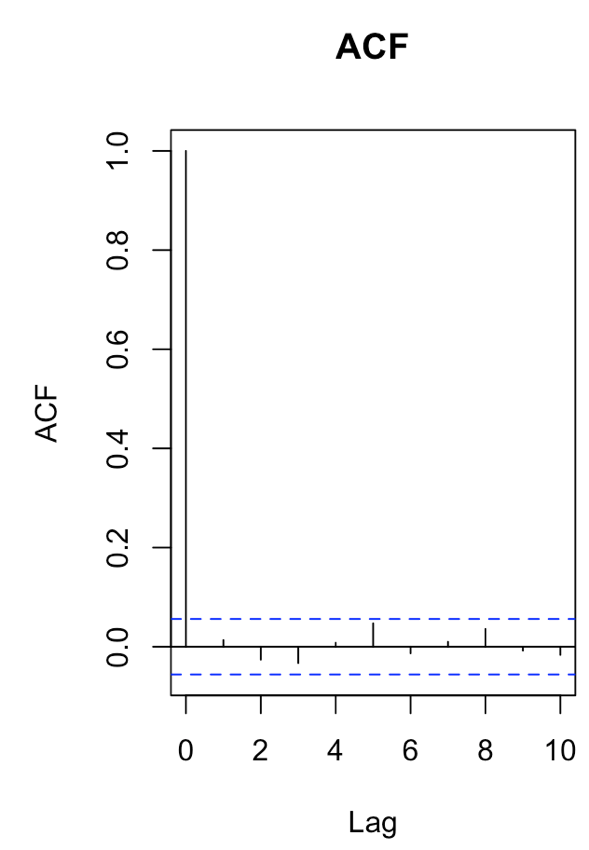

draw(m_number, select=c(1,42))

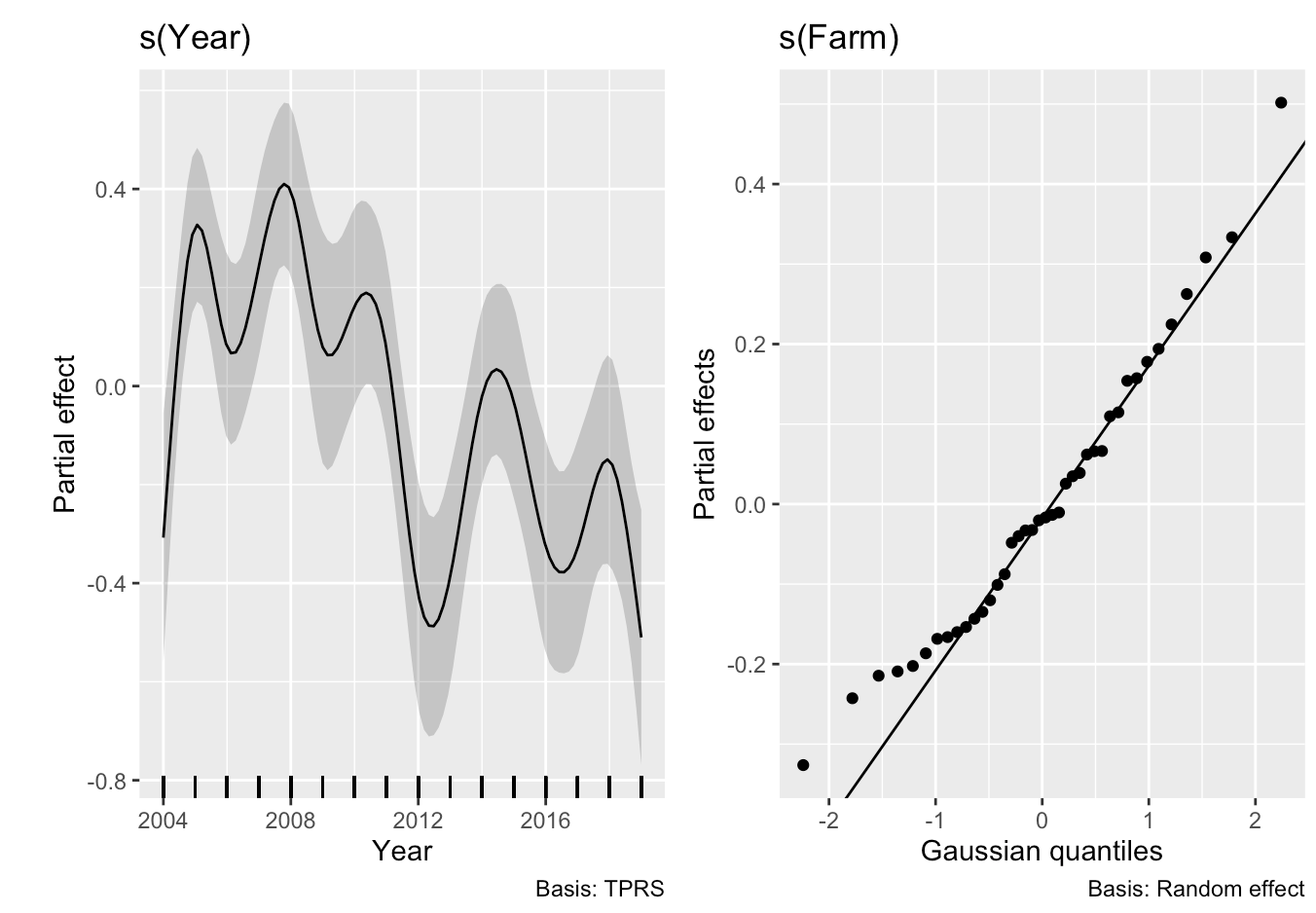

### **Supplementary Material 4 - Geographical distribution of *Protocalliphora* spp. in the study system**

This supplementary figure illustrates the geographic distribution of three *Protocalliphora* species, namely *P. bennetti*, *P. sialia*, and *P. metallica,* across the farms included in our study system. Species proportions were calculated by pooling all individuals of each species recorded on each farm across all years of sampling, thereby neglecting temporal variation. Despite some inter-farm variation, all three species were detected across most of the study area. *Protocalliphora sialia* was by far the most dominant species overall, comprising the majority of records on most farms. This figure provides a spatial overview of *Protocalliphora* community structure within the studied agroecosystem.

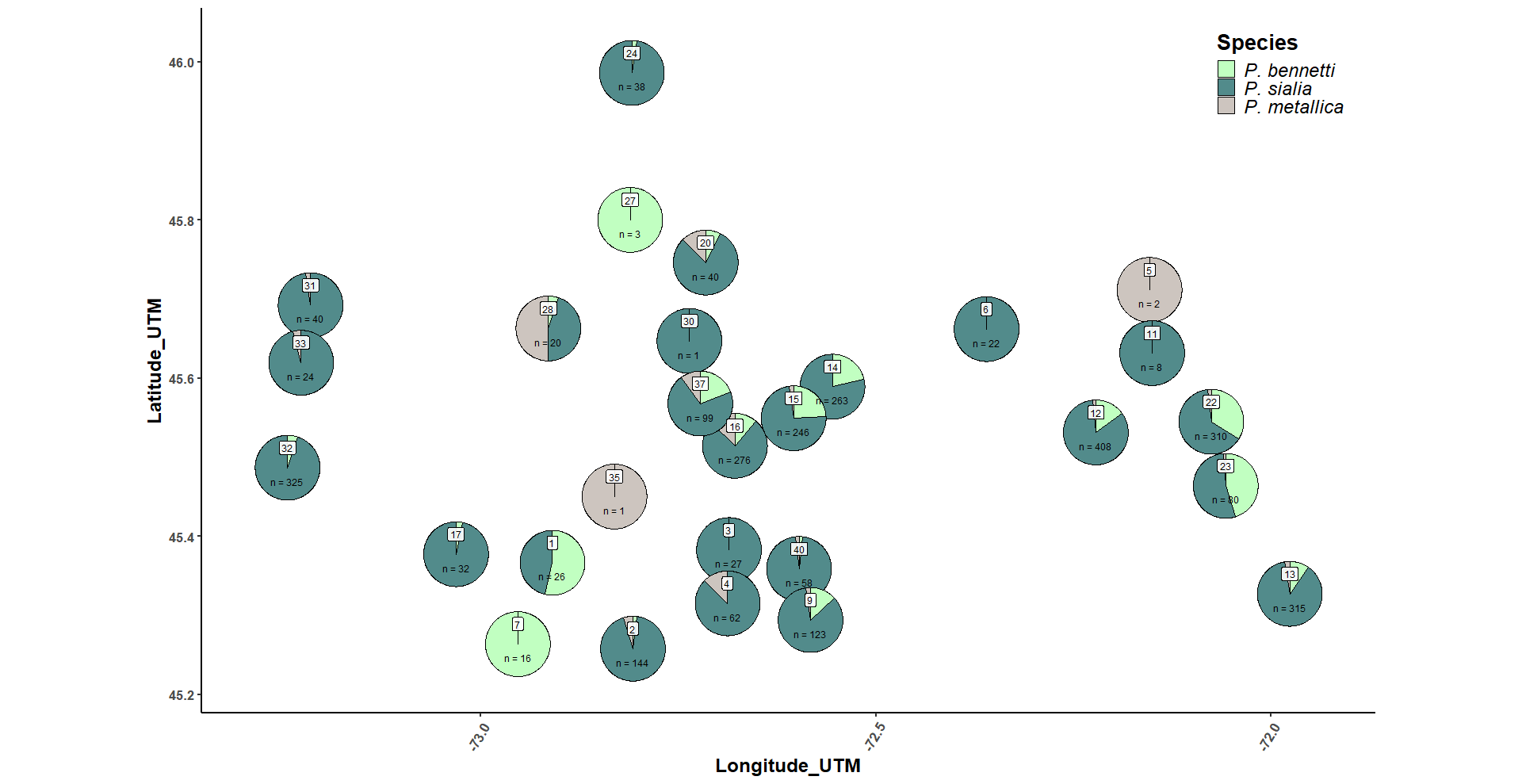
**Figure S4**. Species distribution between *Protocalliphora* *bennetti*, *metallica* or *sialia*, according to the location of the different farms from our study system where specimens were subsampled. Farms were located in southern Québec, Canada and covered 10,200-km2. between 2004 and 2019.

### **Supplementary Material 5 - Interannual variation in infested nests**

This supplementary table reports the annual proportion of Tree Swallow (*Tachycineta bicolor*) nests infested by *Protocalliphora* spp. on each monitored farm over the 16-year study period. For each farm-year combination, the infestation proportion was calculated by dividing the number of infested nests by the total number of occupied nests. Rows represent individual farms, and columns correspond to years. While the exact sample size per farm-year combination is not shown in the table, it ranged from 1 to 10 occupied nests. All nests included in this dataset were sampled and examined for ectoparasite presence. This table provides the raw values that underpin the data presented in Figure 1A of the main manuscript and offers a detailed view of spatial and temporal variation in *Protocalliphora* spp. prevalence across the agricultural landscape.

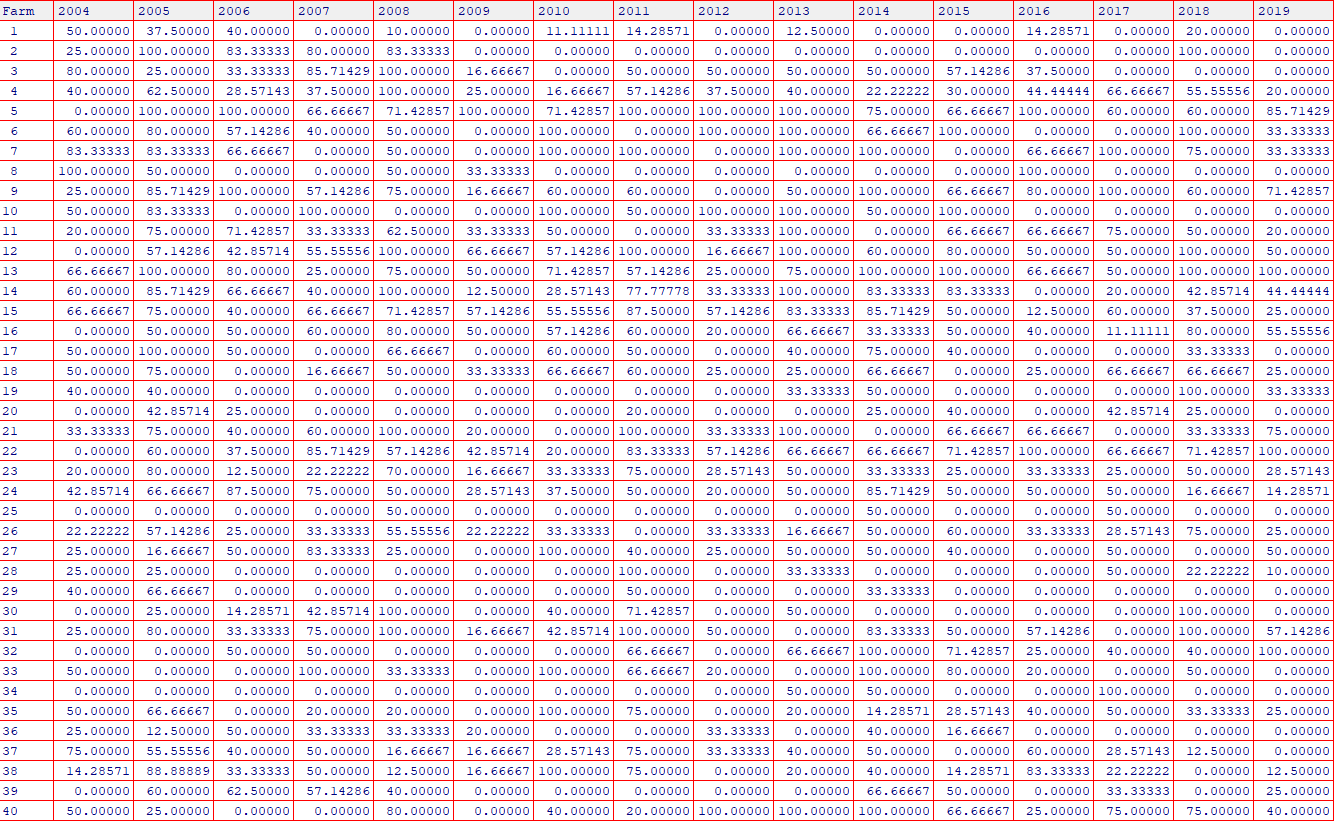
**Table S5**. Proportion of Tree Swallow nests infested by *Protocalliphora* spp. in a given farm according to the year. Number of sampled and examined nests in a farm varies between 1 and 10 per farm per year.
